# Integrated germline and somatic features reveal divergent immune pathways driving ICB response

**DOI:** 10.1101/2024.01.12.575430

**Authors:** Timothy Sears, Meghana Pagadala, Andrea Castro, Ko-han Lee, JungHo Kong, Kairi Tanaka, Scott Lippman, Maurizio Zanetti, Hannah Carter

## Abstract

Immune Checkpoint Blockade (ICB) has revolutionized cancer treatment, however mechanisms determining patient response remain poorly understood. Here we used machine learning to predict ICB response from germline and somatic biomarkers and interpreted the learned model to uncover putative mechanisms driving superior outcomes. Patients with higher T follicular helper infiltrates were robust to defects in the class-I Major Histocompatibility Complex (MHC-I). Further investigation uncovered different ICB responses in MHC-I versus MHC-II neoantigen reliant tumors across patients. Despite similar response rates, MHC-II reliant responses were associated with significantly longer durable clinical benefit (Discovery: Median OS=63.6 vs. 34.5 months P=0.0074; Validation: Median OS=37.5 vs. 33.1 months, P=0.040). Characteristics of the tumor immune microenvironment reflected MHC neoantigen reliance, and analysis of immune checkpoints revealed LAG3 as a potential target in MHC-II but not MHC-I reliant responses. This study highlights the value of interpretable machine learning models in elucidating the biological basis of therapy responses.

**Statement of Significance:** Immune checkpoint blockade works only in a fraction of patients for reasons that are still not fully understood. Our study reveals heterogeneity in the immune responses of ICB responders that correlates with characteristics of the neoantigen landscape. This heterogeneity is accompanied by differences in the duration of clinical benefit as well as by differences as to which immune checkpoint gene serves as a biomarker of ICB response. These findings suggest possible new strategies for improving ICB responses.

**Highlights:**

- We used machine learning to study ICB response across 708 patients from 8 studies across 3 tumor types (melanoma, RCC, and NSCLC).
- Combining germline and somatic features improves prediction of ICB response
- Interactions between germline and somatic features reveal mechanisms contributing to ICB sensitivity.
- MHC-I vs. MHC-II reliance implicates LAG3 as a prognostic biomarker in the context of CD4 T cell driven responses.
- MHC-II neoantigen reliant responses provide superior durable clinical benefit in response to ICB.

## Introduction

The development of immune checkpoint blockade (ICB) drugs has shifted the cancer treatment paradigm, offering unprecedented hope for patients who once faced limited therapeutic options^1,2^. The remarkable successes of ICB, leading to complete remissions in some patients with advanced cancers, have propelled this approach to the forefront of modern oncology^3^. ICB is now a standard treatment in some tumor types, however a substantial proportion of patients still fail to benefit and are needlessly subjected to side effects and costs^4–6^. Despite several landmark studies on biomarkers for immunotherapy response^7–9^ selection of patients who would effectively respond to immunotherapy remains a challenge^10^.

Thus far, biomarkers have focused on measured characteristics of the tumor or the tumor immune microenvironment. Current FDA approved biomarkers include tumor mutation burden (TMB), microsatellite instability (MSI) status, and immunohistochemical staining of the tumor microenvironment (TME) to quantify PD-L1 positivity^11^. However, these predictors of response are imperfect and their application in clinical settings is not straightforward^12^. Many more sophisticated measures of ICB response have also been proposed, including the potential immunogenicity of somatic mutations in the tumor^13,14^, measures of immunoediting such as the ratio of Nonsynonymous/Synonymous mutations of the immunopeptidome^15^, evidence of impaired antigen presentation quantified from somatic copy number loss and mutation of major histocompatibility complex (MHC) genes^16–18^, and tumor clone phylogeny estimates as a proxy for intratumoral heterogeneity^19^. Anagnostou et. al.^20^ successfully integrate somatic features such as these to predict ICB response using machine learning models with superior accuracy, suggesting non-linear predictive models may capture additional biological complexity. These approaches show promise to improve over current FDA based measures, though performance gains have generally been modest.

More recent work has uncovered a role for germline genetic variation in influencing the characteristics of the tumor immune microenvironment and ICB response. Although gold standard whole exome sequencing (WES) methods require a matched normal tissue as a background panel for somatic mutation detection^21^, patient germline variation has largely been ignored in the development of predictive ICB modeling, even though germline variation has a considerable effect on adaptive immune traits^22–24^. We reasoned that while individually, common variants often have only a weak influence on traits, the sum of these variations could have a large impact on the tumor immune microenvironment (TIME) as suggested by a study where common germline variants were found to predict ICB responses independent of somatic biomarkers^25^. With the exception of some rare germline variants, cancer often arises from mutagenic processes independent of host germline genetics^26^.

In this study, we developed a machine learning framework that integrates both somatic and germline features into a unified model that aims to maximize the identification of patients who may benefit from ICB therapy. We used XGBoost for the model architecture as it has shown strong performance on limited training data, allows for non-linear interactions among features, and is interpretable in that individual feature contributions to predictive performance can be quantified^27–29^. A composite model using all features demonstrated superior performance across multiple independent test sets relative to predictors trained on germline or somatic features alone. Analysis of the composite model revealed feature interactions that contributed to model performance, the strongest of which occurred between MHC class-I (MHC-I) damage and a germline variant associated with increased T follicular helper cell infiltration. Further investigation of this interaction suggested an MHC-I independent mechanism of ICB response associated with the MHC class-II (MHC-II) CD4 T cell axis in some patients. Grouping ICB responders by response type showed more durable ICB responses in the MHC-II driven response axis. For the 34% of patients with RNA expression data, we also investigated characteristics of the tumor immune microenvironment such as checkpoint expression, T-cell infiltration, and tertiary lymphoid structure (TLS) signatures^30–32^. Overall, our results support the notion that nonlinear models using somatic and germline features together to predict ICB outcome permit us to formulate new hypotheses about biological mechanisms underlying the diversity of clinical responses to ICB.

## Results

### Design and evaluation of a machine learning framework to predict ICB response

Paired tumor/normal whole exome sequencing (WES) data were obtained for eight independent ICB studies encompassing a range of tissue types and treatments across a total of 708 patients^33–40^. Seven of these were used for machine learning, including feature selection, model training and independent validation (**Fig. 1**), and the eighth (Liu *et. al.)* was added later to validate the translational potential of biological findings. We first assembled a set of germline and somatic features that can be extracted from WES data and that have previously been reported to predict ICB response (**Supplementary Table S1**). Germline SNPs associated with the tumor immune microenvironment (TIME) and ICB response from Pagadala *et. al.*^25^ were further harmonized and aggregated at the gene level into numerical scores for their respective gene, here termed eQTL-scores (**Supplementary Fig. S1,** Methods). SNPs associated with immune infiltration levels were encoded at the single SNP level instead (**Supplementary Table S1**). Somatic features from several impactful ICB response prediction studies were generated for each cohort, including tumor mutational burden (TMB)^41^, dN/dS of the immunopeptidome (ImmunoEditing)^15^, damage of MHC-I alleles^16,17^, and somatic mutation of genes in the antigen presentation pathway^42^. Clinical features available for all data sets included patient age and sex^43^. To train models to predict ICB response, we used a two-stage machine learning approach entailing feature selection followed by model training (**Fig. 1**). We first reduced the number of features via recursive feature selection (RFE) using the Cristescu et. al.^36^ cohort before training an XGBoost^44^ classifier to predict ICB response as class labels (see Methods). XGBoost is a tree-based ensemble method that generates a continuous probability score, here scaled to range between 0-10. We combined three similar anti-PD-1/anti-PD-L1/anti-CTLA4 treated melanoma cohorts (Hugo et. al.^34^, Riaz et. al.^33^, and Snyder et. al.^35^) into a single training set, and evaluated the potential of the classifier to generalize by applying it separately to three heterogenous independent test cohorts: Van Allen (anti-CTLA4 treated melanoma)^40^, Rizvi (antiPD-1 treated NSCLC)^38^, and Miao (anti-PD-1 or anti-PDL1^39^ some also with anti-CTLA4 treated RCC). We compared models that relied only on germline features, only on somatic features, or on a combination of both (referred to as the composite model). We termed the scores produced by these models the immune checkpoint (IC) Index.

**Figure 1:**
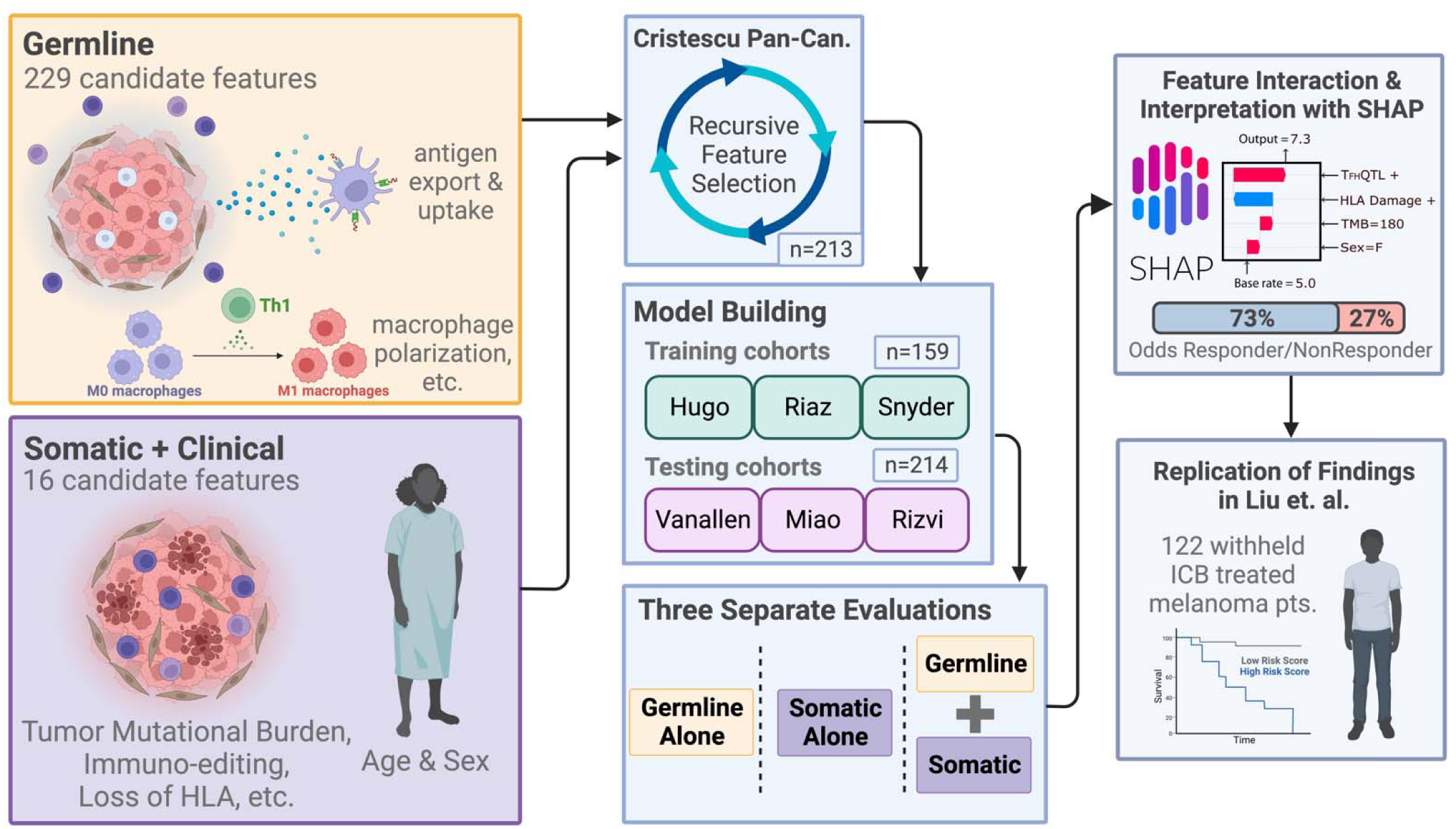
Combining germline and somatic/clinical features into a machine learning framework. Germline and somatic features were assembled from the literature and computed from WES data for 7 ICB studies. Prior to training ICB response predictive models, features and clinical covariates were subjected to recursive feature elimination using the Cristescu study cohort^36^. Models predicting ICB response were then trained on the selected features using three combined studies^33–35^ as the training set. Performance of the trained model was evaluated separately on 3 independent cohorts^38–40^. Models trained only on germline features, only on somatic features and the combination were compared. Feature contributions to the trained model were further investigated to develop biological hypotheses that were reproduced in the Liu cohort^37^.

After recurrent feature elimination (RFE), we retained 24 germline features to train the germline model (**Supplementary Fig. S2A**), including 23 germline eQTL-scores representing genes involved in antigen processing/presentation (*ERAP2*^45^*, ERAP1*^46^, VAMP8^47^), immune signaling (*FCGR2B*^48^*, PDCD1*^49^*, CTSS, CTSW*), and DNA replication (*DHFR*^50^*, TREX1*^51^) and a SNP associated with T-follicular helper cell infiltration^52^ (T_FH_QTL), which was strongly and consistently associated with response across all cohorts (**Fig. 2A**). RFE for the somatic only model selected 13 features derived from clinical and tumor genomic data (**Supplementary Fig. S2B**), including tumor mutational burden (TMB)^12^, clonality-aware derivatives of TMB such as Intra-tumoral Heterogeneity (ITH), and Fraction of TMB subclonal^53,54^, as well as DNA based T cell infiltration estimates^55^, and measures of immune evasion (ImmunoEditing, Immune Escape, MHC-I Damage, Antigen Presentation Pathway Damage)^15–17,56^. RFE for the composite model selected 24 features, 18 (75%) of which were germline eQTL-scores and 6 (25%) of which were somatic features (**Supplementary Fig. S2C**). Considered independently, only a minority of these features showed a significant association with ICB response, and while the direction of effects generally agreed, there was variability across datasets (**Fig. 2A**). Feature associations with ICB response were more similar across melanoma cohorts than other tumor types (**Fig. 2B**). While TMB and clonal TMB features passed recursive feature elimination in the somatic only model (**Supplementary Fig. 3A-C**), they were eliminated in the composite model which instead utilized Fraction of TMB subclonal and ITH– features that are anticorrelated and correlated with TMB respectively, (**Supplementary Fig. S3D-E**; Fraction of TMB subclonal: R=-0.22, P=5.9e-08; ITH: R=0.2, P=1.6e-06). The cIC-Index produced by the trained model remained somewhat correlated with TMB (**Supplementary Fig. S4A**; R=0.2, P=0.0035) even though TMB was not directly incorporated as a feature. The somatic IC-Index (sIC-Index) had an unsurprisingly high correlation with TMB (**Supplementary Fig. S4B**; R=0.46, P=7.5e-13), while the germline IC-Index (gIC-Index) was completely uncorrelated with TMB (**Supplementary Fig. S4C**; R=-0.059, P=0.39).

**Figure 2:**
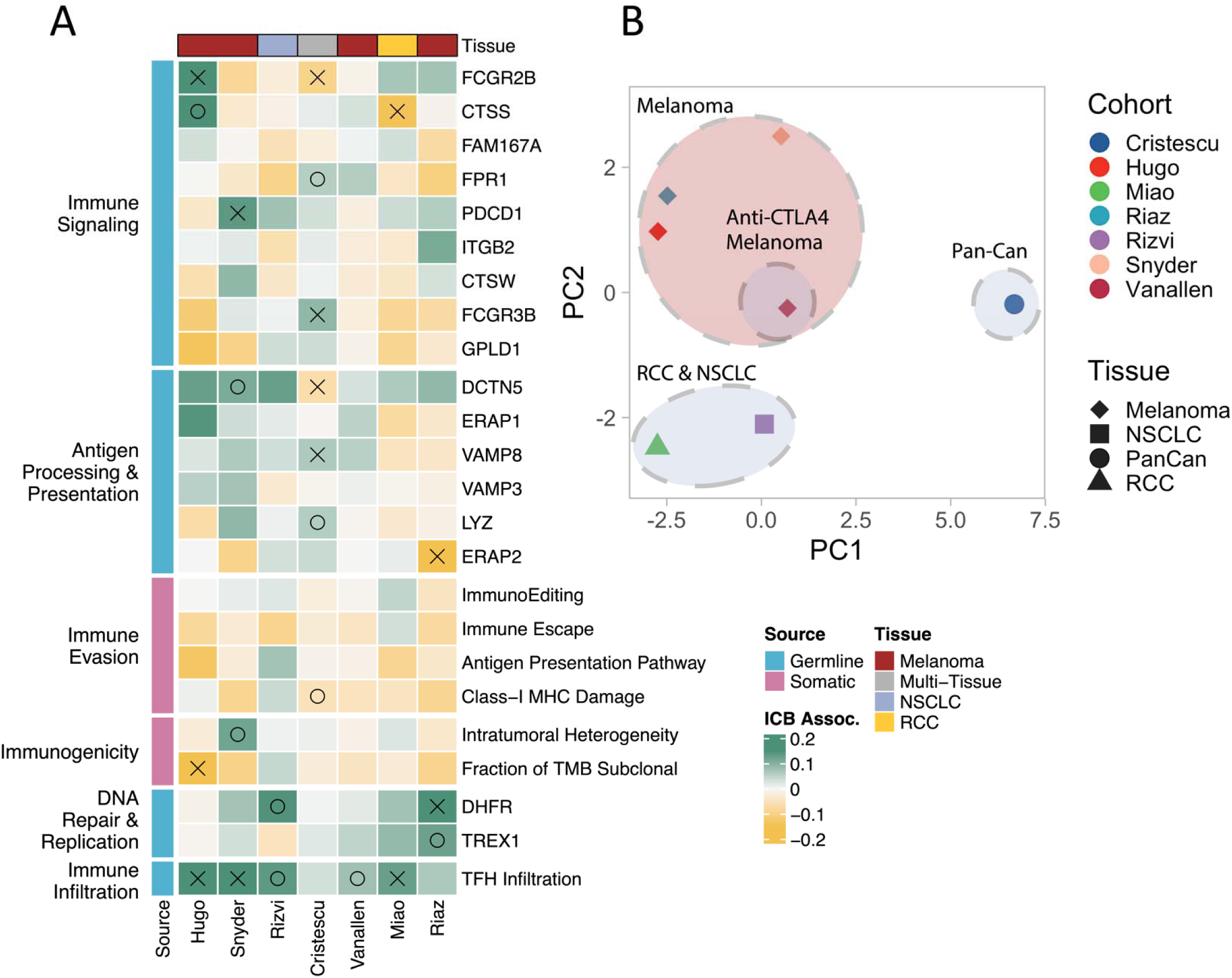
Linear ICB response associations of features passing recursive feature selection. **(A)** Features from germline and somatic models passing secondary RFE on the Cristescu pancancer cohort are shown grouped by cohort and biological impact. Coefficients quantifying associations with ICB response calculated from generalized linear model analysis accounting for age and sex are shown. Green coloring indicates a feature that was more associated with a positive ICB response, yellow indicates association with a negative response. Associations with P<=0.05 are marked with an X, P<=0.1 are marked with O. **(B)** PCA of test statistics for each cohort resulting from linear feature associations.

After training XGBoost models on the selected features using the combined training set, we compared the performance of each model on the three independent test sets. While all three models could distinguish between responders and non-responders, the composite IC Index (cIC-Index) showed the best performance, resulting in the largest mean shift in score distributions between responders and non-responders (**Fig. 3A**), the highest Cliff’s delta between responders and non-responders (**Fig. 3B**), and the highest area under the receiver operating characteristic curve (ROC AUC; **Fig. 3C**). Improvements in ROC AUC from approximately 0.7 to 0.8 were observed in the Van Allen^40^ and Rizvi^38^ studies, but more modest improvements were observed in Miao^39^, possibly due to the vastly different TIME landscape of renal cell carcinomas compared to melanomas^57^. Progression-free survival (PFS) of the highest tertile of cIC-Index scores was significantly higher than the lowest tertile in Kaplan-Meier analysis (**Fig. 3D**, p<0.0001) and the cIC-Index was more predictive of PFS in a Cox proportional hazards analysis using age, sex and tumor type as covariates (see Methods), with a more extreme hazard ratio and more significant p-value relative to germline and somatic only models (**Fig 3E**). Compared to germline and somatic only models, the cIC-Index resulted in an increased positive predictive value (PPV) (**Fig. 3F**, P=0.0012, P=2e-04) while negative predictive power was not significantly different (**Fig. 3G**). Interestingly, gIC-Index and sIC-Index scores were completely uncorrelated with each other, suggesting that these sources of data capture orthogonal information (**Fig. 3H**, R=0.042, P=0.54), helping to explain the improved performance of the composite model. The cIC-Index also outperformed baseline ICB response predictors including TMB, Age, Gender, and checkpoint expression (**Supplementary Fig. 4E-I**)

**Figure 3:**
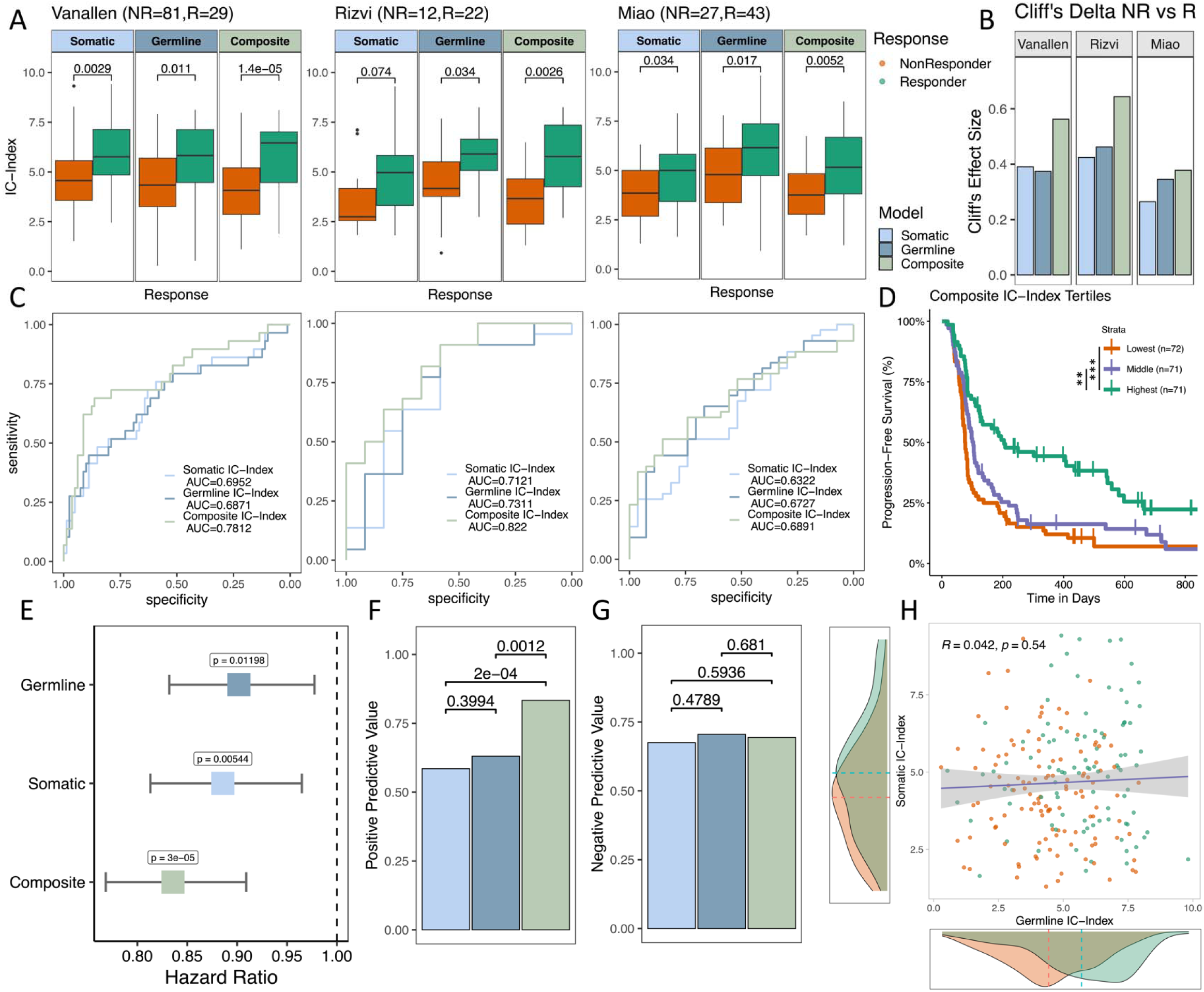
Evaluation of germline, somatic, and composite models across testing cohorts: **(A)** Boxplots of testing cohorts where IC-Index scores are compared via Mann-Whitney U tests. **(B)** Effect size (Cliff’s D) between IC-Index of responders vs nonresponders of each model and cohort **(C)** ROC curves of each model’s performance across testing cohorts with AUC. **(D)** Kaplan-Meier plots of composite IC-Index score tertiles. **(E)** Hazard ratio of IC index for each model from a multivariable Cox PH analysis of progression-free survival across aggregated testing cohorts accounting for age, sex, and cohort. **(F)** Positive predictive value of each model aggregated across all cohorts. **(G)** Negative predictive value of each model aggregated across all cohorts. **(H)** Correlation of germline IC-Index and somatic IC-Index, with somatic and germline model density plots of IC-Index for responders vs nonresponders.

### Impact of Tumor Immune Microenvironment on ICB Response Prediction

Next, we compared the cIC-index to characteristics of the TIME that can be obtained from RNA sequencing data, which were available for (72/214) 34% of test set patients. Several such measures, including effector CD8+ T cell infiltrates^55,58^, joint B and CD4+ T cell levels potentially indicative of tertiary lymphoid structure (TLS) formation^32,59,60^, and target checkpoint expression (PD-L1/CTLA4)^61,62^, have been previously correlated with ICB response. We evaluated CD8+ T cell infiltration levels with CIBERSORTx^63^, a digital cytometry tool that estimates immune cell fractions. To model TLS, we used the gene signature developed by Cabrita et al.^59^ as a proxy for TLS formation. Somewhat surprisingly, patients split by high versus low cIC-Index (cIC-Index>=5) generally had similar TIME infiltration levels in all three categories (**Supplementary Fig. S5A-C**). In converse, the TIME was significantly different between true positives and false positives, where patients who were predicted to respond (cIC-Index>=5) failed to respond and often had an immune-cold TIME, characterized by lower overall levels of immune infiltrates^64^ (**Fig. 4A**). This relationship was strongest in the checkpoint therapy target (CTLA4 for Van Allen et. al., PD-L1 for Miao et. al., Methods) (P=0.0081) and TLS formation TIME categories (TLS gene signature P=0.017), with CD8+ T cells showing near significant association (P=0.055). These results imply that high cIC-Index patients with favorable germline and somatic biomarkers can nonetheless fail to respond to ICB due to a poorly infiltrated TIME.

**Figure 4:**
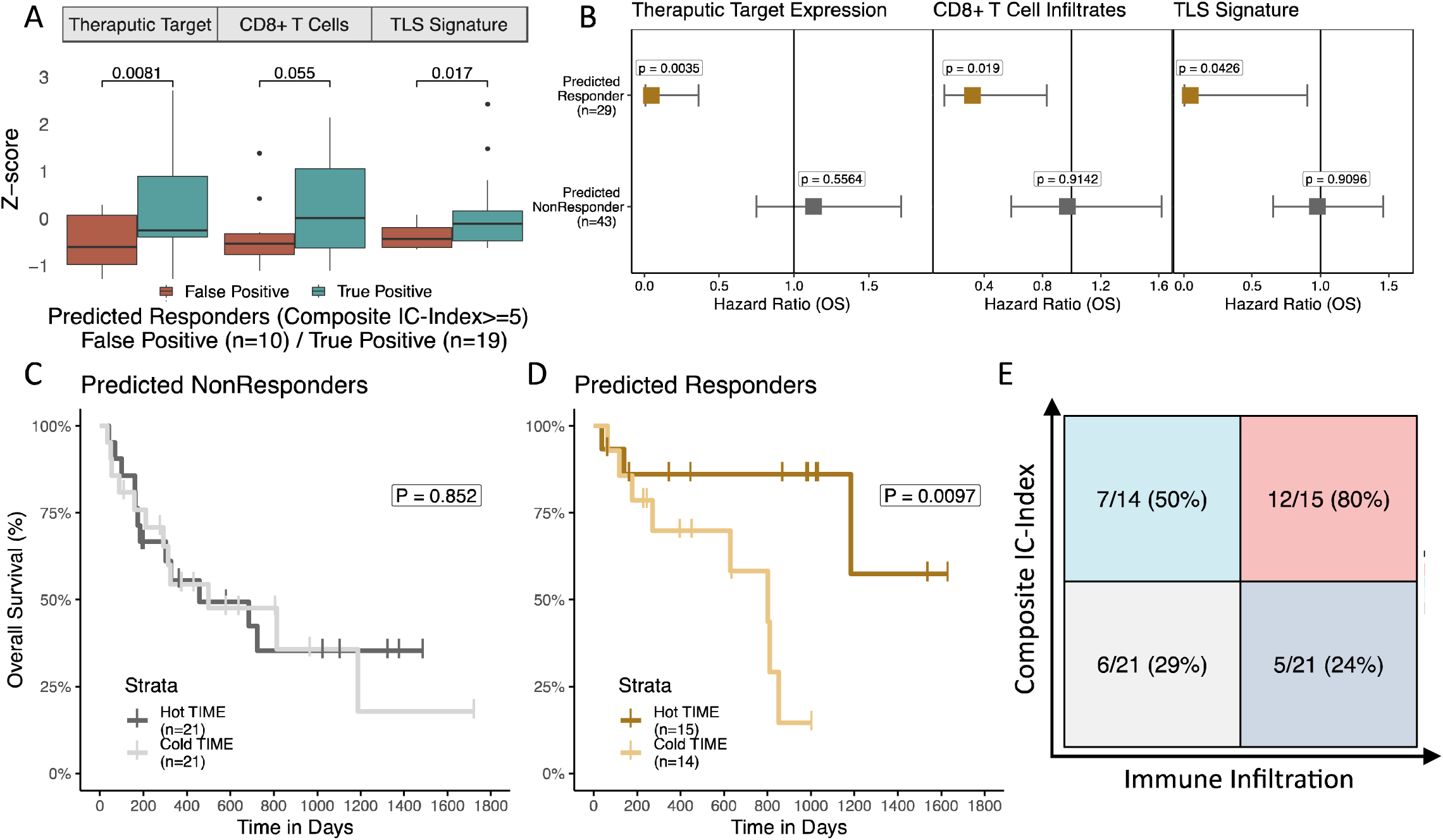
Immune-infiltrated TIME and high cIC-Index scores are synergistic: **(A)** Boxplots of expression-based immune measures from combined test samples with RNA sequencing. IC-Index score distributions are compared with Mann-Whitney U tests. **(B)** Hazard plots of primary TIME biomarkers stratified by IC-Index score. **(D)** Kaplan-Meier curves of predicted nonresponders (IC-Index <5) stratified by TIME biomarker score. **(E)** Kaplan-Meier curves of predicted responders (IC-Index >5) stratified by TIME biomarker score. **(G)** Confusion matrix of IC-Index (cutoff IC-Index >5) and TIME Score (cutoff TIME Score above median).

We also investigated whether an immune hot TIME could rescue patients with low somatic and germline potential for response. Using a Cox proportional hazards model adjusted for age, sex, and data set, we found that each of the TIME infiltration estimates (checkpoint target: P=0.0035, CD8 T cells: P=0.019, TLS formation: P=0.043) was significantly associated with improved overall survival in high cIC-Index patients only, whereas low IC-Index patients failed to significantly benefit from an immune hot TIME (**Fig. 4B**). These results are mirrored in Kaplan-Meier plots of high and low cIC-Index patients (**Fig. 4C-D**) stratified by level of TIME infiltration (Methods). High cIC-Index patients benefit from an above median TIME (P=0.0097), while low cIC-Index patients do not (P=0.852). These findings are consistent with previous studies indicating that immunogenic tumors respond at greater rates when there is high CD8+ T cell infiltration, but that high CD8+ T cell infiltration alone is not sufficient for high rates of ICB response^36^. Interestingly, while high cIC-Index scores yielded the strongest relationship with higher immune infiltration, we found this synergy was primarily driven by germline factors rather than somatic ones (**Supplementary Fig. S5D-E**). Our analyses suggest that cIC-Index scores may be useful as general estimates of immunogenicity and could be used as additional indicators of when a patient could benefit from ICB beyond TIME profiling.

### Non-linear feature interactions reveal alternative mechanisms of ICB response

In order to better understand how selected germline and somatic features contribute to model performance, we analyzed feature importance using SHAP values^65^, a game theory approach to improve the interpretation of the machine learning model. We noted differences in feature rankings particularly for ERAP1, MHC-I damage, and Immunoediting, between XGBoost and linear models suggesting the presence of interactivity effects (**Supplementary Fig. S6**). Thus, we evaluated both individual feature contributions and pairwise interactions between features. SHAP analysis revealed several key feature interactions (**Fig. 5A**), the strongest of which was between the somatic MHC-I damage, *i.e*., the cumulative MHC-I damage from somatic mutation and loss of heterozygosity (see Methods), and the T_FH_QTL. We further examined this interaction in terms of ICB response rates between categories (**Fig. 5B**) and observed higher rates of response when the T_FH_QTL is present (P=1.7e-05 T_FH_QTL vs class-I MHC damage, P=0.0063 T_FH_QTL vs neither), even when the potentially negative effect of MHC-I damage is present (P=1.0, T_FH_QTL vs both). Because rates of ICB response are unaffected by MHC-I damage in patients carrying the T_FH_QTL (**Fig. 5B**), we hypothesized that this SNP may promote immune responses upon ICB treatment that do not rely on MHC-I based antigen presentation, suggesting instead a role for MHC-II driven mechanism of response.

**Figure 5:**
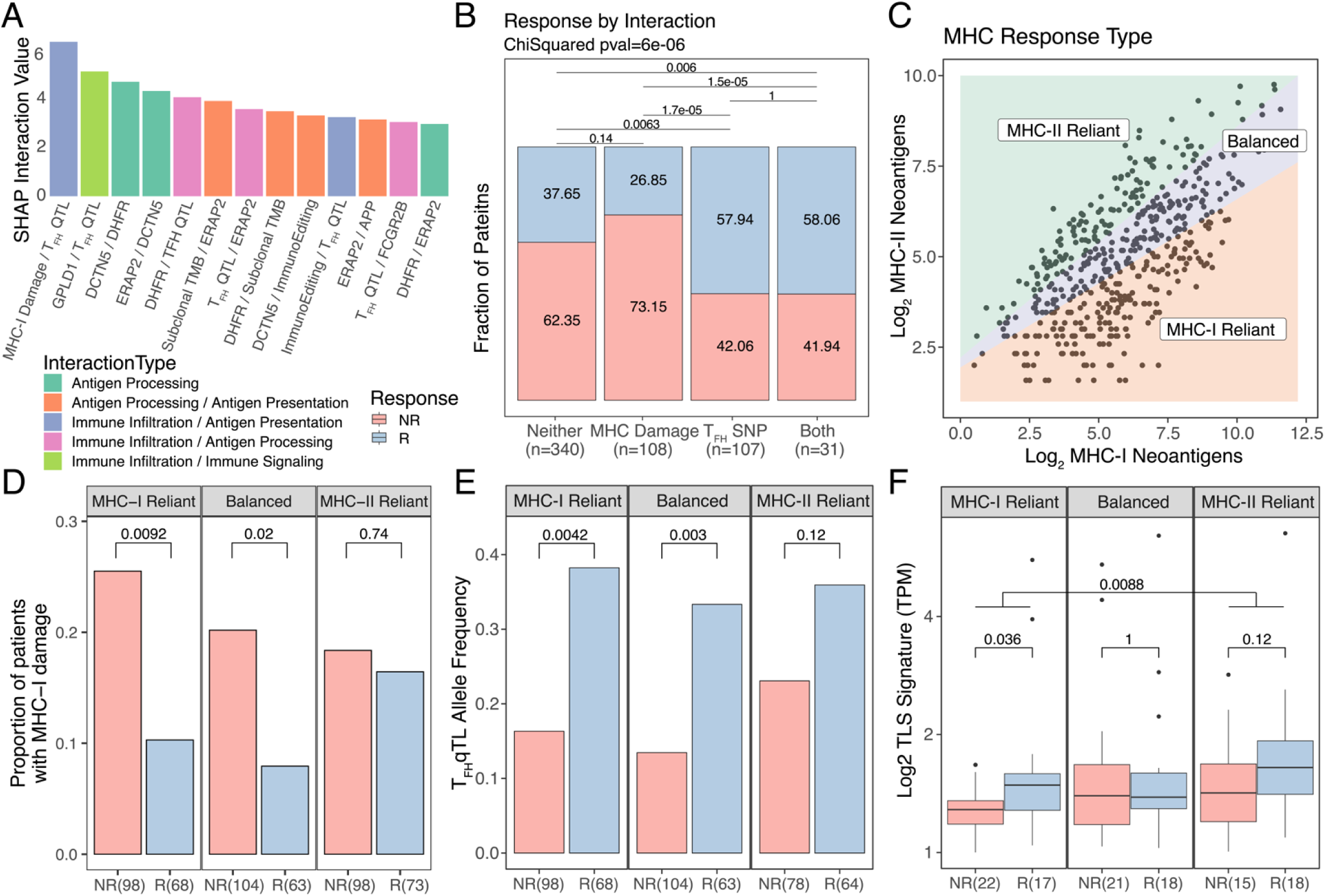
Non-linear feature interaction analysis reveals differences in response mechanisms. **(A)** Shap interaction scores for the top 12 interacting feature pairs. InteractionType categories reflect main biological relevance of constituent features. **(B)** Bar plot of percent responder by interaction category for the MHC damage and T_FH_ SNP interaction category. **(C)** Schematic of MHC based response groupings. **(D)** Proportion of patients with MHC-I damage split by MHC response categories and response status. **(E)** Allelic fraction of the T_FH_QTL faceted by MHC response categories and split by response status. **(F)** TLS signature expression split by MHC response categories and response status.

To further investigate this idea, we grouped tumors in the dataset according to whether somatic mutations were more prevalently presented by MHC-I or MHC-II molecules, suggesting the potential for reliance of immune responses on particular MHC pathways of neoantigen presentation. First, we calculated PHBR scores^66,67^ (see Methods) for each nonsynonymous mutation in all patients. PHBR scores are mutation-centric scores that seek to summarize whether any peptides overlapping the mutated site will be presented by any of an individual’s HLA alleles. Patients with at least three mutations passing PHBR thresholds for both class-I and class-II MHCs were then split into groups termed MHC-I reliant, MHC-II reliant, or balanced based on the ratio of these class specific neoantigens (**Fig 5C**), with reliant referring to an immune response potentially dependent on MHC-I vs MHC-II presented neoantigens. Among MHC-I reliant patients, unsurprisingly we noted a significantly higher level of MHC-I damage in nonresponders vs responders (P=0.0092, **Fig. 5D**) reflecting the notion that an MHC-I reliant response depends on the integrity of the MHC-I and associated antigen presentation pathway. While Balanced patients demonstrated an intermediate disparity in MHC-I damage between nonresponders vs responders (P=0.02), this was not the case in MHC-II reliant patients (P=0.74). Overall ICB response rates between these two groups were not significantly different (**Supplementary Fig. S7A**).

Next we sought to understand how MHC-reliance could modify potential to benefit from the T_FH_QTL. We reasoned that the most extreme cases of MHC-II reliance would be those that also had defects in the MHC-I antigen presentation pathway. This group comprised 83% of MHC-II reliant tumors (154/171), so we focused further analyses on this aspect (see Methods). We found a significant difference in the frequency of the T_FH_QTL between responders vs nonresponders in the MHC-I reliant and balanced categories (P=0.0042, P=0.003, **Fig. 5E**), but not in the solely MHC-II reliant category (P=0.12). This is somewhat mirrored in the subset of patients with tumor immune infiltration estimates available, where T_FH_ cell estimates were higher in MHC-I reliant responders vs nonresponders (P=0.03, **Supplementary Fig. S7B**) but not in the balanced or MHC-II reliant responders vs nonresponders (P=0.48, P=0.5). It is possible that MHC-I reliant responders benefit from an increased infiltration by T_FH_ cells, TLS formation and associated helper effects that are important to maintain the function and precursor frequency of CD8 T cells^68–73^. Indeed, TLS have been shown to enhance ICB response in melanoma^59,60^. Conversely, MHC-II reliant patients may receive less benefit from additional T_FH_ cell infiltration because their neoantigen landscape is already predisposed towards the formation of TLSs. Indeed, we found that MHC-I reliant responders had higher TLS gene signature expression than nonresponders (P=0.036, **Fig. 5F**), yet this difference was not significant in MHC-II reliant patients (P=0.12, **Fig. 5F**). MHC-II reliant patients in general had a higher level of TLS gene signature expression than MHC-I reliant patients (P=0.0088, **Fig. 5F**), which is not altogether surprising given that TLS formation is more closely associated with the MHC-II / CD4+ T cell axis^74–76^. These initial observations point to the possibility that mechanistically divergent immune responses yield ICB response based on how effectively neoantigens engage each MHC pathway.

### MHC reliance groupings are related with survival and mechanism of immune evasion

We next sought to understand the clinical implications of differential MHC reliance. To validate our findings, we performed identical analyses on an additional independent ICB treated cohort (n=77) with paired transcriptomic data (Liu et. al.^37^) and compared to our original set of seven cohorts referred to as the discovery set. We first investigated effects of MHC reliance grouping on the composition of the tumor immune microenvironment (see Methods). Interestingly, we observed that CD4/CD8 T cell ratios mirrored MHC reliance in responders, with higher ratios being observed in MHC-II reliant tumors (**Fig. 6A**, P=0.0057 Discovery; P=0.025 Validation). However, no such a difference was found in nonresponders. We applied an identical methodology to immune-infiltrated ^77^ ICB-naive, tissue matched cancer samples from TCGA (see Methods) and found a powerful protective effect by the CD4/CD8 ratio in TCGA MHC-II reliant patients (**Supplementary Fig. S8A**, HR=-0.76, P=0.0069), but a significantly adverse effect of that same ratio in TCGA MHC-I reliant patients (**Supplementary Fig. S8B**, HR=0.59, P=0.0352). These data support a benefit to having some level of concordance between CD4/CD8+ T cell infiltration and MHC-II/MHC-I neoantigen ratios. To investigate differences in response dynamics between CD4+ and CD8+ T cell mediated responses, we compared the survival of responders MHC-II vs MHC-I reliant groups. Despite nonsignificant differences in response rates, MHC-II reliant responders had a significantly longer overall survival in both discovery and validation cohorts (**Fig. 6B-C**, discovery P=0.0073; validation P=0.0398), consistent with reports that CD4+ T cell based immune responses are tumor autonomous and therefore more difficult to evade in the long term^67,78,79^.

**Figure 6:**
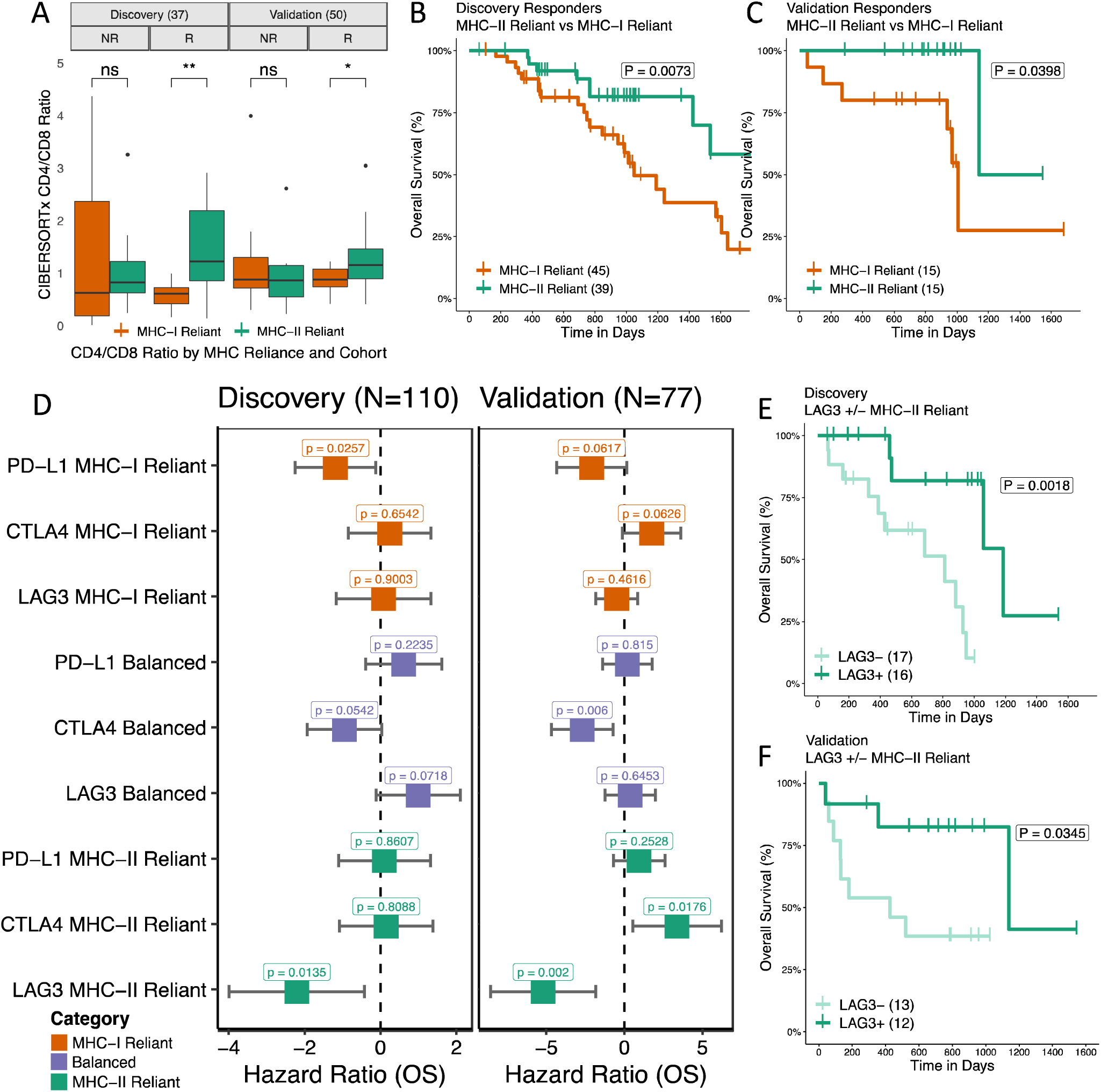
TIME, survival duration, and checkpoint expression impact differs by MHC reliance status. **(A)** CIBERSORTx CD4/CD8 T cell infiltration estimate ratios split by MHC Reliance category and response status for both discovery and validation cohorts. **(B-C)** Kaplan-Meier curves of overall survival of responders only in MHC-II reliant patients vs MHC-I reliant patients for discovery and validation cohorts. Schematic of MHC based response groupings. **(D-E)** Hazard ratios of key checkpoint molecule expression levels colored by MHC reliance status. Age, sex, and cohort specific covariates are included in multivariate analysis. **(F-G)** Kaplan-Meier curves of LAG3 above/below median expression in MHC-II reliant patients in discovery and validation cohorts.

Finally, we wanted to know if differences in MHC reliance could translate to differences in pathways of immune evasion. Immune checkpoints are commonly overexpressed to suppress an active immune response. Currently, of the many the checkpoints identified in the TME only PD-L1 positivity in tumor sections is approved as a biomarker of ICB response, albeit its predictive value is modest ^61^. To investigate whether differences might exist as to which checkpoints correlate with a beneficial anti-tumor immune response under different MHC reliance conditions, we evaluated the relationship between expression of individual checkpoint genes and PFS post-ICB treatment by univariable Cox PH analysis. We focused on checkpoint genes with antibody inhibitors undergoing clinical trials (PD-L1, CTLA4, LAG3, TIGIT, TIM3, IDO1, and OX40)^80^. When split by MHC reliance grouping, higher LAG3 expression was associated with benefit from immune checkpoint blockade in the MHC II reliant group (**Supplementary Fig. S9A-D**). To adjust for potentially confounding effects of the correlated expression of canonical^81,82^ immune checkpoint genes, we performed a multivariable analysis centered on LAG3, PD-L1, and CTLA4 (see Methods). We found that high PD-L1 expression was generally associated with longer survival post ICB treatment in MHC-I reliant patients (**Fig. 6D-E**, discovery P=0.026; validation P=0.062), CTLA4 expression with longer survival in balanced patients (**Fig. 6D-E**, discovery P=0.054; validation P=0.006), and LAG3 with longer survival in MHC-II reliant patients (**Fig. 6D-E**, discovery P=0.014; validation P=0.002). There was no association of checkpoint gene expression with MHC reliance category (**Supplementary Fig. S10A-B**). Among MHC-II reliant patients, higher expression of LAG3 was associated with significantly longer overall survival in both discovery and validation cohorts (**Fig. 6F-G**, discovery P=0.0018; validation P=0.0345). LAG3 is thought to play a prominent role in CD4+ T cell regulation and may be a primary marker of activation^83,84^. Our results may, therefore, reflect a key role for LAG3 as a mediator of CD4+ T cell based response to ICB therapy.

## Discussion

Immune checkpoint blockade has emerged as a potent anti-cancer therapy, however the fraction of patients that benefit from treatment remains disappointingly low. To improve the success of ICB, it is of the utmost importance to understand which factors govern the potential to respond via the immune system. Here we used a machine learning framework to study somatic and germline biomarkers of response to ICB in human cohorts. We were able to extract both feature types from paired tumor-normal whole exome sequencing data across eight ICB-treated human studies. Germline immune eQTL biomarkers, while relatively new, show promise to capture complementary information from somatic features, and XGBoost models trained to predict a composite IC index (cIC-Index) using both feature types performed better at predicting ICB response across different tumor types. When we interrogated patients with additional available RNAseq data, we found that the survival benefit of an immune hot microenvironment was contingent upon having a high cIC-Index score, that there was no response in patients with a low cIC-Index score, and that this was driven, surprisingly, by germline features. This supports the notion that heritable differences in immune cell function determine the effectiveness of an immune response once immune cells have reached the tumor. Furthermore, patients with a high cIC-Index score who failed to respond often had a “cold” tumor immune microenvironment. This suggests that transcriptomic profiling might be useful as a supplemental prognostic tool of ICB response in high cIC-index patients, and that the cIC-Index score serves as a general proxy for clinical response to the immune invigorating effect of ICB.

To gain further insight as to how various biomarkers relate to ICB response potential, we used state-of-the-art techniques for interpreting machine learning models, and studied important features and feature interactions that drove model predictions. Surprisingly, the strongest interaction involved an interplay between a SNP associated with increased T-follicular helper cell infiltration (T_FH_QTL) and MHC-I damage. Specifically, we observed a beneficial effect of the T_FH_QTL on rates of response, independent of the deleterious effect of MHC-I damage. T follicular helper (T_FH_) cells are the specialized subset of CD4^+^ T cells that help B cells produce antibodies in germinal centers (GC)^76^. T_FH_ cells are normally located in secondary lymphoid organs at close distance with B cells^85^. However, there is increasing evidence that T_FH_ cells are part of tertiary lymphoid structures (TLS), intra-tumor organized clusters of immune cells including B and T cells and dendritic cells (DCs) mimicking germinal centers in secondary lymphoid organs^76,86^. TLS are an increasingly common finding in cancer, and are linked with better prognosis^87,88^; increased infiltration by T_FH_ cells and TLS formation are a source of helper factors beneficial to both CD8+ and CD4+ T cells. Indeed, the number of TLS distinguishes ICB responders from non-responders^32,60^.

MHC-I damage on cancer cells inherently hampers the cytotoxic function of CD8+ T cells, yielding low response rates. Surprisingly, we found that response rates were rescued when patients had both the T_FH_QTL and MHC-I damage, suggesting that rescue mechanisms of ICB response may be shifted towards MHC-II mediated immunity (MHC-II reliance). Using individual level information about the ratio of neoantigens with binding affinity for MHC-I and MHC-II, we were able to allocate patients to either a MHC-I or MHC-II reliant group. That these groupings may initiate and sustain differential immune mechanisms in response to ICB is strengthened by the observation that MHC-II reliance promotes higher infiltration of CD4 T cells and more durable clinical responses to ICB, potentially reflecting a direct effect on long-term memory CD4+ T cell responses. In contrast, MHC-I reliant responses, which are centered on CD8+ T cells, are possibly more transient in the absence of CD4 T cell help^69^.

When we examined the association of pre-treatment checkpoint gene expression levels with ICB response, which was predominantly anti-PD1/anti-PD-L1 treatment in the cohorts studied, we found that PD-L1 expression was associated with better ICB response in MHC-I reliant patients but not in MHC-II reliant patients while the reverse was true for LAG3. In patients where immune evasion is mediated by over-expression of PD-1/PD-L1, anti-PD1/anti-PD-L1 therapies can be remarkably effective^89^. LAG3 on the other hand has MHC-II as its major ligand^84^ and it is widely regarded as a negative regulator of CD4+ T cell activation^90^. Higher expression of LAG3 could therefore indicate an effective ongoing MHC-II reliant anti-tumor response pre-ICB treatment. In our analysis, LAG3+ patients had better survival in the MHC-II reliant group, suggesting that MHC-II driven immunity can support an effective response to anti-PD1/anti-PD-L1, and that this could potentially be further amplified by an anti-LAG3 therapy. However, the lack of association of PD-L1 expression with response in the MHC-II reliant group seems to suggest a mechanism independent of alleviating PD-L1 based repression of CD8 T cells. A similar phenomenon has been observed in microsatellite instable colorectal cancers with B2M loss that paradoxically remain among the best responders to anti-PD1/anti-PD-L1 therapy^91^. It is intriguing to think that anti-PD1/anti-PD-L1 can be beneficial even if PD-L1 is not highly expressed or the class-I antigen presentation machinery is not functional. Recent data show that LAG3 also associates with the T cell receptor (TCR)-CD3 complex in both CD4^+^ and CD8^+^ T cells in the absence of binding to MHC-II, causing the dissociation of the tyrosine kinase Lck from the CD4 or CD8 co-receptors and loss of co-receptor-TCR signaling during T cell activation^92^. Our finding that LAG3 facilitates the CD4 T cell response during ICB treatment could be explained by the fact that both LAG3 and ICB target the proximal signaling of the T-cell receptor^93^, even though the reasons this creates an advantage in MHC-II reliant patients remains unclear. Perhaps this reflects the fact that the adult peripheral repertoire is richer in CD4+ than in CD8+ T cells. This bias may also explain the observation that cancer patients vaccinated with neoantigens have a propensity to generate CD4 T cell responses^94^.

The other implication is that the utility of each of these checkpoint genes as biomarkers of ICB response may be highly context dependent. PD-L1 expression was not associated with ICB response in MHC-II reliant patients responding via a CD4+ T cell axis of adaptive immunity. This could explain in part why PD-L1 positivity is a surprisingly poor general predictor of response rates^95^. Future efforts to refine biomarkers of ICB response could attempt to leverage widely available germline information as well as understand the context of a patient’s MHC reliance status.

Our study was subject to some limitations. Publicly available ICB treated cohorts with DNA sequencing data remain relatively limited and RNA data is even more so. Larger feature selection and training cohorts could further improve model performance. Future studies could incorporate additional biomarkers, for example genotypes associated with adverse immune events, such as rs16906115 affecting *IL7*^96^, that could lead to early stopping of therapy, or copy number alterations affecting key immune loci^97,98^. We also limited our features to those extractable from paired tumor-normal WES as tumor DNA to mirror what is more commonly available in real world settings. While the germline derived features in the composite model are straightforward to compute once bioinformatic infrastructure is in place, the variety and complexity of the somatic features may be more challenging to implement in the clinic. MHC Reliance groupings were based solely on single nucleotide variants. Future versions of our PHBR pipeline will include support for frameshift and stoploss variants, which may be more impactful in an immunogenicity context. Most ICB response classification approaches eliminate difficult to classify Stable Disease (SD) patients from their studies—despite the fact that these patients benefit from increased survival from ICB treatment. We chose to include these patients as responders to maximize potential clinical benefit, at the cost of increasing the complexity of our classification task. Finally, while our classifier–which was trained on melanoma patients–showed some ability to generalize to other tumor types, especially non-small cell lung cancer, it may ultimately be essential to train and study tumor-type specific models.

## Conclusion

Investigation of the factors that determine ICB response in cancer patients is providing key insights into mechanisms that drive superior response. This study provides further evidence that CD4 T cell responses engaged by MHC-II antigen presentation are a critical component of superior immune responses, and points to an alignment of checkpoint-based evasion with the particular immune cell types dominating the response. This sets the stage for future strategies to optimize selection of checkpoint therapies from characteristics of the patient tumor and immune system.

## Methods

### ICB Data Sets

Raw FASTQ files were obtained using SRA toolkit v2.9.6-1-ubuntu64 for the following immune checkpoint trials: Hugo et al. 2016 (SRA accession: SRP090294, SRP067938; Cancer: melanoma), Van Allen et al. (SRA accession: SRP011540, Cancer: melanoma), Miao et al. (SRA accession: SRP128156, Cancer: clear cell renal carcinoma), Riaz et al. (SRA accession: SRP095809, SRP094781; Cancer: melanoma), Rizvi et al. (SRA accession: SRP064805, Cancer: non-small cell lung cancer), Snyder et al. (SRA accession: SRP072934, Cancer: melanoma), Liu et. al. (SRA accession: Cancer: melanoma), and Cristescu et. al. (SRA accession: PRJNA449580, Cancer: Melanoma, HSNCC, Urothelial). Only pre-treatment samples were utilized in this study. Across cohorts, a total of 708 ICB treated patients were evaluated in this study.

### Data processing

FASTQ files were processed via an identical bioinformatics pipeline. **DNA:** Genomic reads were aligned to UCSC hg19 coordinates using BWA v0.7.17-r1188. Reads were sorted by SAMTOOLS v0.1.19, marked for duplicates with Picard Tools v2.12.3 and recalibrated with GATK v3.8-1-0. Germline variants were called from sorted BAM files using DeepVariant v0.10.0-gpu. Somatic variants were obtained through the following additional steps. Aligned tumor/normal BAM files were submitted to standard Mutect2 somatic variant calling using GATK-4.1.3.0. First, BAM file formats were standardized using GATK-4.1.3.0 AddorReplaceReadGroups, then GATK-4.1.3.0 Mutect2 was used to call somatic variants using default settings (including the presence of a matched normal), the gnomAD v3.1 raw sites background SNP panel, and the Twist Exome Target bed file to limit variant calling to exonic regions. Potential somatic variants were filtered using GATK-4.1.3.0 FilterMutectCalls and only mutations with a filter flag of “PASS” were kept for subsequent analysis. Somatic mutations were further filtered to retain only those with a DNA allelic fraction > 5%. The resulting VCF files were annotated by VEP using cache version 102_GRCh37 and default settings. **RNA:** Where available, RNA FASTQ/BAM files were downloaded for 33 RCC and 240 melanoma patients. BAM files were converted to FASTQ using bam2fq. Unpaired reads were removed using fastq pair. Paired reads were aligned with STAR v2.4.1d to GRCh37 reference alignment. RSEM v1.2.21 was used for transcript quantification. Raw transcript counts were corrected for cohort specific batch effects using ComBat before being transformed into TPM values.

### Feature construction

#### Germline features

A set of 1084 tumor immune microenvironment (TIME) associated SNPs were sourced from Pagadala et. al^25^. These SNPs were demonstrated to have significant associations with immune related functions in TCGA, and were successfully used to develop an earlier germline ICB response prediction model. Next we filtered for SNPs present with a MAF > 0.05 in all studies, leaving 598 SNPs to run METAL^99^ analysis with ICB response in the three training cohorts. METAL analysis calculates a single P-value for each SNP across the three training cohorts (Hugo et. al., Riaz et. al., and Snyder et. al.) and indicates the direction of effect for each cohort. SNPs with an FDR < 0.25 and showing full agreement of direction of impact were included, resulting in 229 SNPs with a nominal ICB association. TCGA and discovery genotype processing was performed in Pagadala et. al. and is described in detail in their methods. For this study, we obtained pre-processed genotype matrices for each of the cohorts examined.

#### Somatic features

##### Tumor mutational burden

Tumor mutational burden (TMB) was defined as the sum of all nonsynonymous somatic coding mutations in each patient’s VCF file, including “protein coding”, “frameshift variant”, and “stop lost” mutations. To adjust for cohort specific effects, TMB was transformed by the intra-cohort z-score before being included in the machine learning model. A similar convention is described in Vokes et. al^100^.

##### Immune Escape

A comprehensive list of immune escape related genes was obtained from Zapata et. al^15^. Somatic mutations with VEP impact annotations of “MODERATE” or “HIGH” were tallied from per patient VCF files. The final Immune Escape mutation counts were divided by each patient’s total TMB to generate a score reflecting disproportionate immune evasion—otherwise the score is highly correlated with TMB.

##### Antigen Presentation Pathway

A list of key antigen presentation pathway related genes was obtained from MSigDB M1062, Reactome Antigen Presentation Folding Assembly and Peptide Loading of Class-I MHC. All HLA genes were removed from this list as they are accounted for with better accuracy by HLA specific tools and summarized in other features. Somatic mutations with an impact of “MODERATE” or “HIGH” were tallied from per patient VCF files. The resulting scores were divided by each patient’s total TMB to generate a score reflecting disproportionate damage to the antigen presentation pathway.

##### IntraTumoral Heterogeneity and Fraction of TMB Subclonal

IntraTumoralHeterogeneity and Fraction of TMB Subclonal both rely on accurate subclonal estimates, which are derived as follows. First, copy number calling was performed using CNVkit v0.9.10. A background panel of normals was constructed for each cohort separately using CNVkit reference to protect against batch effects. CNVkit batch was used to call copy number changes with each respective cohort’s matched background panel. We next used PureCN v2.6.4 (run via singularity image) with CNVkit derived .cnr and .seg files, and Mutect2 derived filtered VCF files to generate purity and ploidy metrics to be used in subsequent subclone estimation. PureCN was run with default settings, repeat regions censored, and a random seed set to 123. Next, PyClone-VI v0.13.1 was run on mutation specific integer copy number estimates derived from CNVkit call (https://cnvkit.readthedocs.io/en/stable/heterogeneity.html) to estimate clonal structure of the tumor. IntraTumoral Heterogeneity (ITH) was defined as the total number of subclones with at least 5 mutations (total range 0-11 subclones). Fraction of TMB Subclonal was calculated by taking the total number of mutations belonging to small subclones (<5 mutations per subclone) and dividing by the total number of mutations for each tumor. This generates an inverse estimate of clonal heterogeneity from ITH.

##### ImmunoEditing

ImmunoEditing evaluates the ratio of nonsynonymous to synonymous mutations (dN/dS) in a tumor as a measure of selection^101^. Immune dN/dS was adapted by Zapata et. al.^15^ in their toolkit SOPRANO (https://github.com/luisgls/SOPRANO) to calculate the ImmunoEditing score for each patient using an hg19 reference and default settings. Essentially, this score derives from calculating dN/dS across all regions of the proteome predicted to bind the set of patient-specific MHC alleles (i.e. displayed for immune surveillance) and ranges from 0 to ∼5 with a score above 1 indicating a higher amount of nonsynonymous mutations to synonymous ones.

##### Class-I MHC Damage

Class-I MHC Damage was defined as the union of POLYSOLVER^16^ and LOHHLA^17^ results. First, Class-I HLA alleles were genotyped via POLYSOLVER (See PHBR pipeline methods). Next, LOHHLA (https://github.com/mskcc/lohhla), originally published in McGranahan et al., is used to identify copy number losses of HLA alleles. Copy number and purity data is provided to the program and summary statistics about HLA copy number losses are generated. A given HLA allele was marked as lost if the Pval_unique of its loss was <=0.05. POLYSOLVER mutation calling (Shukla et. al.) was used to generate somatic mutation calls of each HLA allele. If an HLA allele was flagged by either of these tools, it was marked as damaged. Alleles were only counted as damaged once even if flagged by both tools. Both programs were provided identical HLA genotypes on a per patient basis.

### Machine learning framework

#### Overview

We built XGBoost classifiers for three predictive tasks: ICB response prediction from germline, somatic and combined features respectively. Models were fit in 2 stages: feature selection, followed by model training and evaluation. First, we conducted Recursive feature elimination (RFE) on an initial array of features using the Cristescu et. al. cohort, then trained classifiers to predict ICB response using Hugo et. al. (34), Riaz et. al. (61) and Snyder et. al. (64) melanoma cohorts. The trained model was then evaluated on 3 test cohorts: Vanallen et. al. (110), Miao et. al. (70) and Rizvi et. al. (34). Biological implication validation was conducted with the Liu et. al. (122) cohort.

#### Recursive Feature Elimination (RFE)

Recursive feature elimination was performed on three feature sets: 229 germline SNPs only, 16 somatic variables only, and both sets combined. The recursive feature elimination model was trained on Cristescu melanoma (89) and tested on Cristescu HNSCC (107) and Cristescu urothelial (17) samples to ensure this step prioritized broadly useful biological features to use in the model training step. The model used for RFE was an XGBoost Random Forest Classifier (python package version 1.6.2) with 20 total estimators and a maximum depth of 8. We used a nonlinear model for feature selection to allow for feature interactions even during the feature selection stage. All possible feature combinations and total model sizes were tested and the mean squared error (MSE) of each was recorded. The model with the lowest MSE was selected, and the features included in that particular model were used for training in stage two. For the 229 germline SNPs, a model with a combination of 54 SNPs yielded the lowest MSE in the RFE cohorts. These 54 SNPs were collapsed into continuous gene level eQTL-scores by measuring the direction of their effect on gene expression in TCGA and orienting alleles such that all SNPs affected gene expression in the same direction (Fig. S1). This resulted in 23 simplified, gene-level continuous scores reflecting the total magnitude of expected change in gene expression (**Supplementary Fig. S1A**). For the composite model, RFE was performed on the set of features prioritized by the initial RFE performed for each data type separately.

#### ICB Response Classifier Training

We trained three different classifiers to predict ICB response, one using only germline features, one using only somatic features and one on the combined feature set (the composite model). Using features passing RFE analysis, XGBoost Random Forest Classifiers were trained on Riaz et. al, Hugo et. al., and Snyder et. al. data sets with 1200 total estimators and a maximum depth of 8. The performance of these models was then evaluated separately on the Vanallen et. al., Rizvi et. al., and Miao et. al. datasets. Aside from feature curation, this process was identical for all models. and a standard random seed was set for all models to ensure reproducibility. For each patient, the XGBoost Random Forest Classifier returns a class prediction probability ranging from 0 to 1, which we refer to as the IC-Index. For visualization purposes, we used sklearn MinMaxScaler to scale these values from 0-10. This process preserves the distribution of scores and therefore does not affect statistical comparisons. For each model and cohort, IC-Index scores were compared between responders and nonresponders using Mann-Whitney U tests. Receiver Operating Characteristic (ROC) plots were constructed using the scaled continuous IC-index results where the outcome label was the response phenotype, and the area under the curve (AUC) was used to summarize overall performance. Test datasets were then pooled for survival analysis via multivariable Cox Proportional Hazards analysis, where the association of IC-Index with progression-free survival was measured alongside covariates of age, sex, and tumor type, using the R packages “survival” and “survminer”^102,103^. Kaplan-Meier curves were constructed using tertile splits of IC-Index scores and P-values of pairwise comparisons between tertiles were computed with log-rank tests. Finally, positive and negative predictive values (PPV and NVP) were computed and compared between each model type using the “DTComPair” package^104^. State of the art ICB response prediction projects from Litchfield et. al., Chowell et. al., and Auslander et. al.^105–107^ have demonstrated remarkable accuracy in validation sets when RECIST stable disease (SD) category patients are included as nonresponders or excluded entirely. These SD patients are particularly difficult to classify due to their ambiguous TIME and somatic biomarker landscape, but still benefit from increased overall survival^108^ and were counted as responders in predictive modeling tasks.

### Evaluation of the tumor immune microenvironment with digital cytometry

The composition of immune infiltrates in the tumor immune microenvironment (TIME) was evaluated by digital cytometry via CIBERSORTx using the LM22 signature matrix with batch correction. The T Cell Infiltration score was constructed from the CIBERSORTx CD8 T Cells score. The general TIME score used in Kaplan-Meier plotting was calculated as the linear combination of Therapeutic Target, T Cell Response, and TLS Formation. CIBERSORTx T follicular helper cell estimates were reused for MHC Reliance analyses to corroborate the effect of the T_FH_QTL. The tertiary lymphoid structure gene expression signature was generated from a set of TLS related genes reported by Cabrita et al and Sautès-Fridman et. al. (CCL19, CCL21, CXCL13, CCR7, CXCR5, SELL, LAMP3, CETP, RBP5, AICDA, BCL6, CCR6, CD79B) using the method put forth in Cabrita et. al. where mean gene expression of key genes upregulated in TLS was calculated. CD4 and CD8 T cell infiltration estimates were calculated using CIBERSORTx, where the CD4/CD8 ratio was defined using “T cells CD4 memory.activated” + “T cells follicular helper” infiltration divided by “T cells CD8” infiltration categories. Only patients in the top two tertiles of CD8 T cell infiltration were included in direct CD4/CD8 ratio comparison analysis to remove patients with zero or very low levels of immune infiltrates.

### SHAP feature importance and feature interactions

Feature importance and interaction within non-linear models were calculated using the SHAP machine learning interpretability suite (https://shap.readthedocs.io/en/latest/). SHAP, which stands for SHapley Additive exPlanations, is a unified approach to explain the output of any machine learning model. It is based on cooperative game theory and the concept of Shapley values. SHAP values assign each feature an importance value for a particular prediction in the context of a specific model. These values allow for nonlinear interactions between features to be accounted for on a per-patient basis, and also allow us to rank pairwise feature interaction by magnitude. Each model was run through the standard SHAP python pipeline and the feature importances were recorded (**Supplementary Fig. S3**). For the composite model, feature interaction analysis was performed as well using the shap_interaction_values function.

### PHBR score pipeline

Originally developed by Marty et. al.^66,67^, the Patient Harmonic-mean Best Rank (PHBR) score is a measure of how well a given neoantigen is presented by the major histocompatibility complex (MHC) based on computationally derived binding affinities between all possible peptides harboring the mutation and a patient’s set of HLA alleles. A detailed description can be found in the original publication^66^. For each patient, all single nucleotide variant mutations were given an MHC-I PHBR score and an MHC-II PHBR score representing presentation by class-I and class-II respectively. A neoantigen was considered to be well presented by MHC-I with a PHBR score <=2, and well presented by MHC-II with a PHBR score <=10^109^. Class-I HLA alleles were called using POLYSOLVER^16^ (v1.0.0) with default parameters, and Class-II HLA alleles were called using HLA-HD^110^ (v1.4.0) with default parameters.

### MHC reliance stratification

Patients were stratified by the ratio of the total number of neoantigens well presented by class-II MHC divided by the total number of neoantigens well presented by class-I MHC. A patient was only considered for analysis if they had at least three mutations well presented by both MHC-I and MHC-II. Neoantigens that were both well presented by both MHC-I and MHC-II were not considered in this ratio. These ratios were divided into tertiles and defined as follows: the lowest tertile was MHC-I reliant, the middle tertile was balanced, and the highest tertile was MHC-II reliant. To select for patients with MHC-II based immune responses, MHC-II reliant patients with no evidence of MHC-I damage or loss of heterozygosity were excluded.

### TCGA immune infiltration analysis

Tissue types matching those from our analysis (melanoma, renal cell carcinoma, non-small cell lung carcinoma, head and neck squamous cell carcinoma, and urothelial/bladder cancer) were pulled from TCGA (LUAD, KIRC, SKCM, HNSC, BLCA, KICH, KIRP, LUSC). Stage II-IV cancers were analyzed to better match our ICB cohorts. Poorly infiltrated tumors were dropped from the analysis to ensure that cancers analyzed from TCGA were at least somewhat infiltrated by lymphocytes. To achieve this, we calculated the ImmunoScore^77^, for all patients, and the bottom tertile (most poorly infiltrated) patients were dropped from the analysis. CD4/CD8 T cell ratios were calculated in an identical manner as the ICB cohorts. Similarly, MHC Reliance groupings were generated identically as in ICB discovery and reliance cohorts.

### Multivariable checkpoint analysis

Five FDA unapproved immune checkpoint genes with ongoing clinical trials were investigated for an association with a particular MHC Reliance group: LAG3, TIM3, TIGIT, OX40, and IDO1. Univariable analysis revealed significant associations with LAG3 in both discovery and validation cohorts, which was subjected to further multivariable analysis accounting for PDL1 and CTLA4 expression. A median expression cutoff was used to create binary high and low expressing groups for each of the checkpoint genes. Age, sex, and tumor type were accounted for during multivariable analysis, as well as prior CTLA4 treatment in the validation cohort, due to a large proportion of patients in Liu *et. al.* having received such treatment. Kaplan-Meier curves were generated using these same binary cutoffs and P-values were calculated using the log-rank test.

## Data Availability

This study relied entirely on published datasets. All datasets were obtained via dbGaP or SRA at the following accessions: Rizvi *et. al.* phs000980.v1.p1; Riaz *et. al.* SRP095809; Miao *et. al.* phs001493.v2.p1; Van Allen *et. al.* phs000452.v2.p1; Hugo *et. al.* SRP067938 and SRP090294; Snyder *et. al.* SRP072934; Cristescu *et. al.* PRJNA449580; Liu *et. al.* phs000452.v3.p1

## Code Availability

Code to reproduce models, analyses and figures can be found at the following Github repository: https://github.com/cartercompbio/MHC_reliance

## Acknowledgements

This work was funded by Mark Foundation Emerging Leader Award #18-022-ELA, NCI grant R01CA269919 and support from NCI grant U24CA248138 to HC. Computational resources were supported by infrastructure grant 2P41GM103504-11. The results shown here are in part based upon data generated by the TCGA Research Network. ICB datasets: For Rizvi *et al.* non-small cell lung cancer immunotherapy analysis, we used dbGaP data from accession phs000980.v1.p1. We thank the members of the Thoracic Oncology Service and the Chan and Wolchok labs at MSKCC for helpful discussions. We thank the Immune Monitoring Core at MSKCC, including L. Caro, R. Ramsawak, and Z. Mu, for exceptional support with processing and banking peripheral blood lymphocytes. We thank P. Worrell and E. Brzostowski for help in identifying tumor specimens for analysis. We thank A. Viale for superb technical assistance. We thank D. Philips, M. van Buuren, and M. Toebes for help performing the combinatorial coding screens. This work was supported by the Geoffrey Beene Cancer Research Center (MDH, NAR, TAC, JDW, AS), the Society for Memorial Sloan Kettering Cancer Center (MDH), Lung Cancer Research Foundation (WL), Frederick Adler Chair Fund (TAC), The One Ball Matt Memorial Golf Tournament (EBG), Queen Wilhelmina Cancer Research Award (TNS), The STARR Foundation (TAC, JDW), the Ludwig Trust (JDW), and a Stand Up To Cancer-Cancer Research Institute Cancer Immunology Translational Cancer Research Grant (JDW, TNS, TAC). Stand Up To Cancer is a program of the Entertainment Industry Foundation administered by the American Association for Cancer Research. For Snyder *et al*. melanoma immunotherapy analysis, we used dbGaP data from accession phs001041.v1.p1. We thank Martin Miller at Memorial Sloan Kettering Cancer Center (MSKCC) for his assistance with the NetMHC server, Agnes Viale and Kety Huberman at the MSKCC Genomics Core, Annamalai Selvakumar and Alice Yeh at the MSKCC HLA typing laboratory for their technical assistance, and John Khoury for assistance in chart review. For Miao *et al.* renal cell carcinoma immunotherapy analysis, we used dbGap data from accession phs001493.v2.p1. This study was supported by an AACR KureIt grant. Hugo *et al.* melanoma samples were acquired from SRA using accession numbers SRP067938 and SRP090294. Riaz *et al.* melanoma samples were acquired from SRA using accession number SRP095809. For Van Allen *et al.* melanoma sample, data was acquired from dbgap accession phs000452.v2.p1. For Liu *et. al.* melanoma validation cohort, data was acquired from dbgap accession phs000452.v3.p1 and supported by the National Human Genome Research Institute (NHGRI) Large Scale Sequencing Program, Grant U54 HG003067 to the Broad Institute (PI, Lander).

## Author Contributions

Concept and study design: T.S., H.C. Data processing, analysis and machine learning: T.S., M.P., A.C., K.T. Supervision: HC. Manuscript writing T.S., S.L., M.Z., H.C., Scientific and editorial feedback: K.L., J.K., S.L., M.Z., H.C.

## SUPPLEMENTARY FIGURES

**Fig S1.**
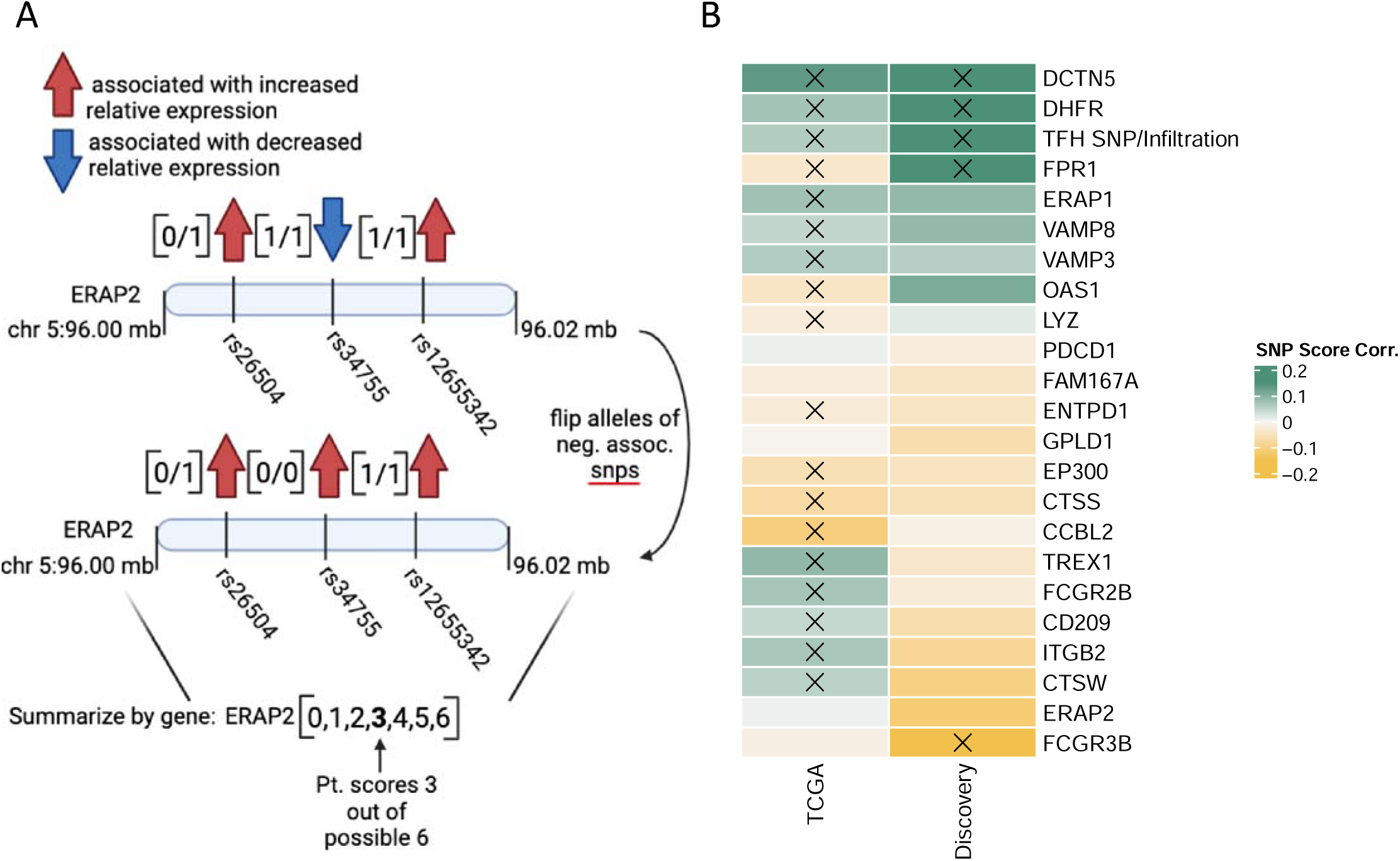
**A.** eQTL-score schematic. SNPs alleles are oriented so that they are all in alignment in terms of direction of effects on gene expression. Number of alleles affecting gene expression are then summed into a continuous score **B.** Pearson correlation of Gene level eQTL-scores in TCGA and Discovery cohorts. P<=0.05 is indicated by an X.

**Fig S2.**
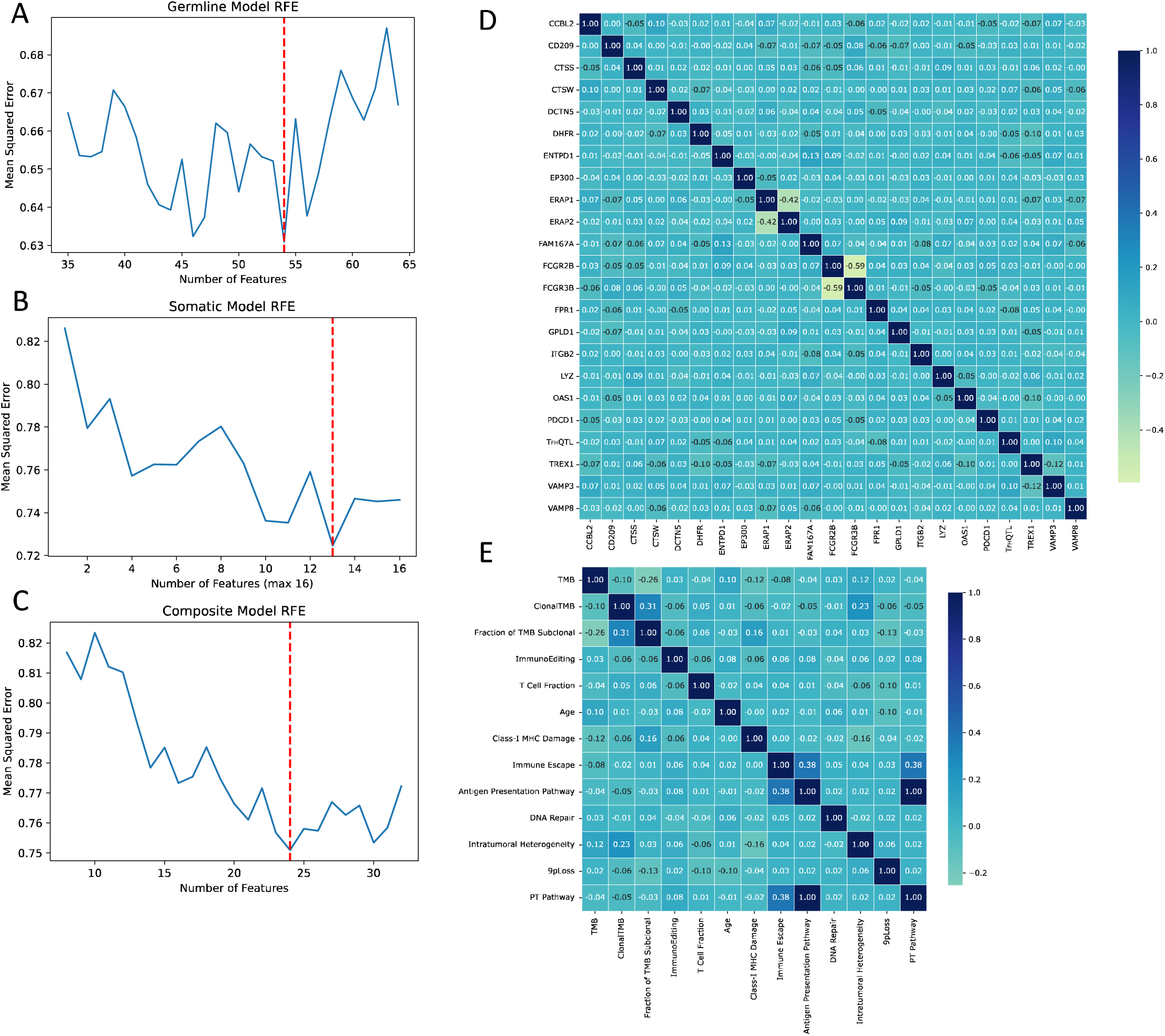
**A-C**. Recursive feature elimination analysis. Mean squared error for each model is shown for a range of total number of features included in a given model. Optimal model size is indicated by a dashed red line. **D-E.** Pearson correlation matrices of features respective to germline and somatic models.

**Fig S3.**
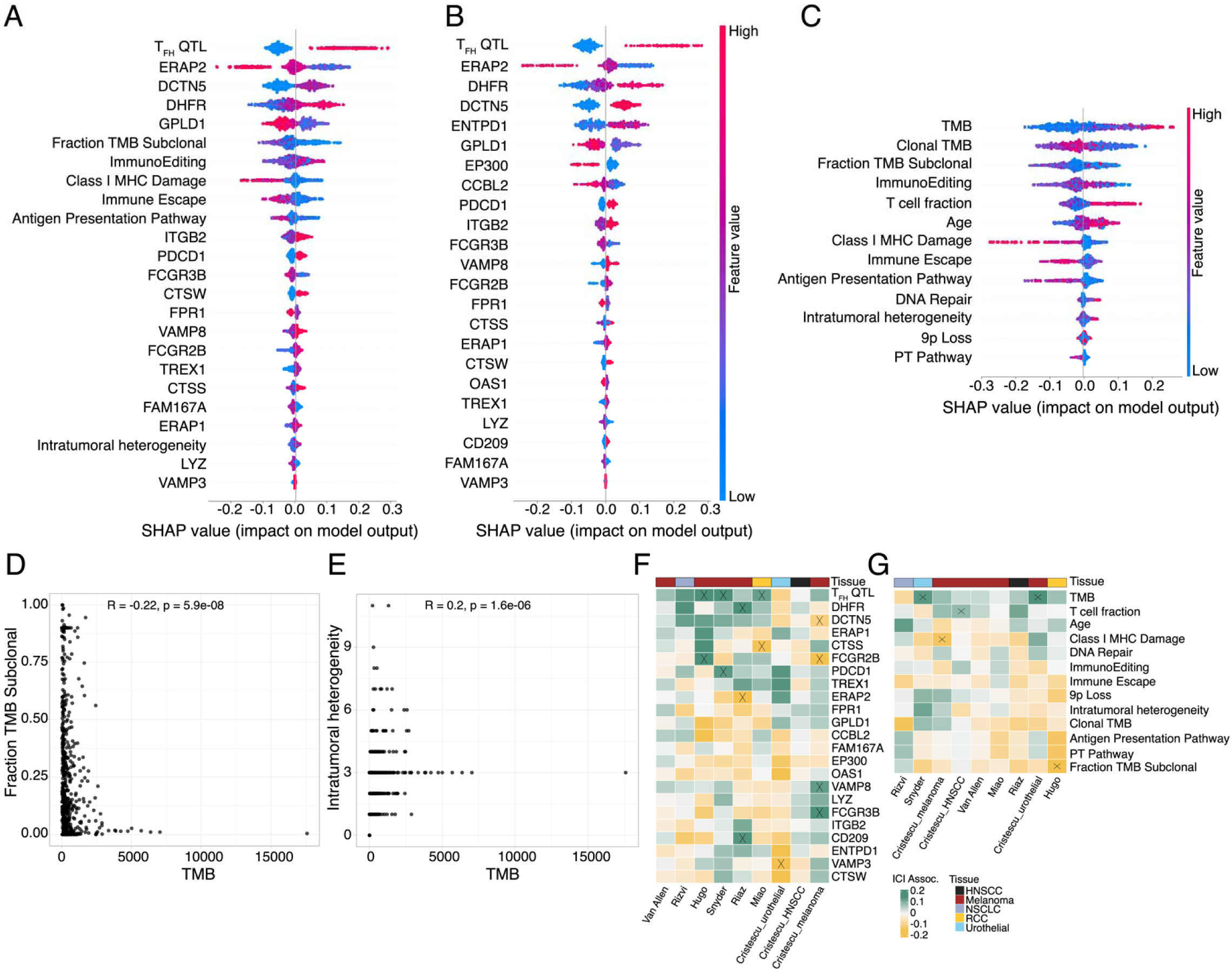
**A-C**. SHAP derived nonlinear feature importance beeswarm plots of germline, somatic, and composite models. **D-E.** Correlation of immunogenicity features with TMB. **F-G.** Coefficients of features from linear regression-based germline and somatic models

**Fig S4.**
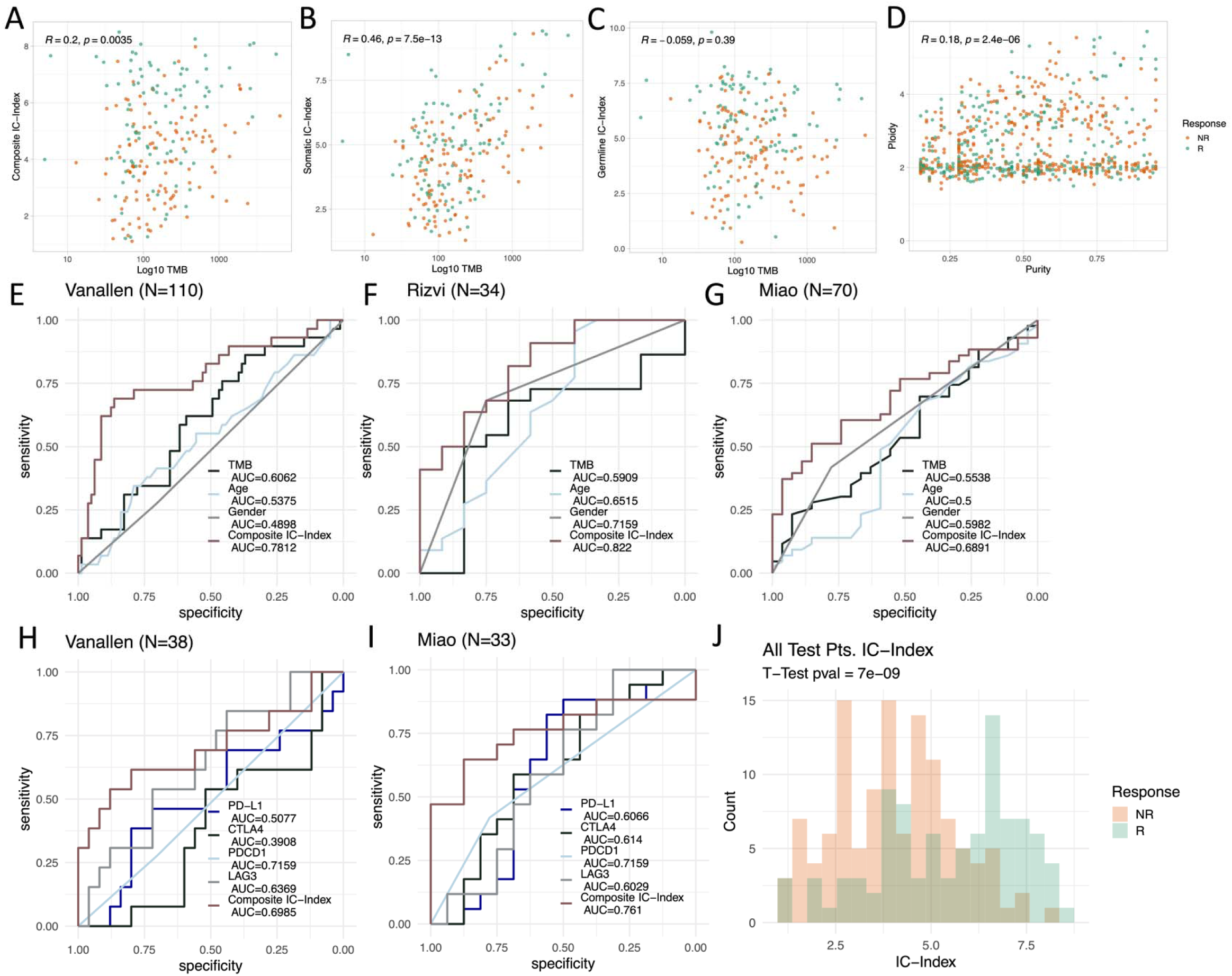
**A-C.** Pearson correlation of three different IC-Index scores with TMB. **D.** Purity vs ploidy across all patients, colored by response with pearson correlation shown. **E-G**. ROC plots comparing the performance of composite IC-Index to TMB and clinical predictors of ICB response. **H-I.** ROC comparing the performance of composite IC-Index to transcriptomic predictors of ICB response **J.** Histogram of composite IC-Index across patients, colored by ICB response status.

**Fig S5.**
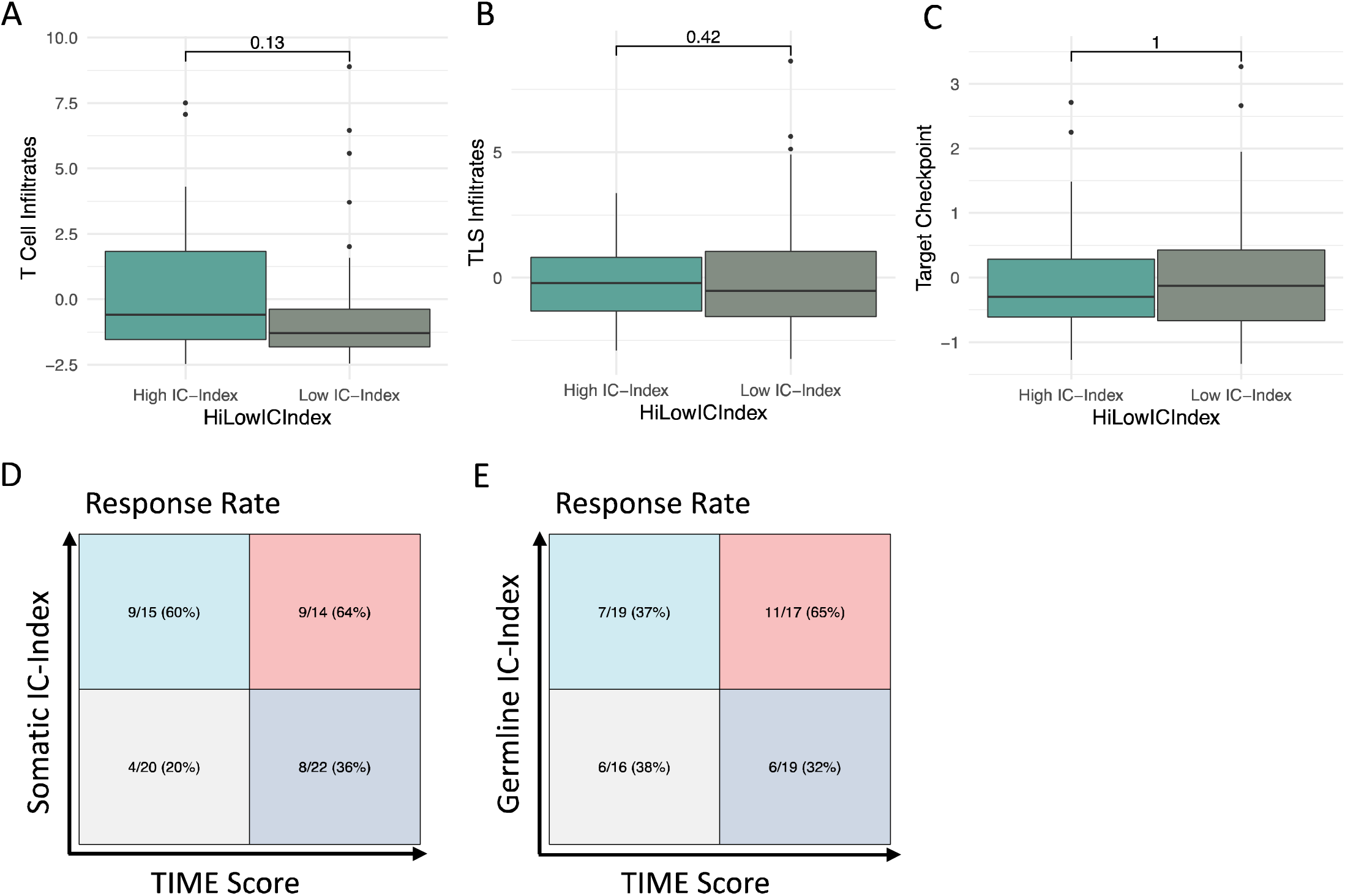
**(A-C)**. Measures of TIME infiltrates stratified by high vs low composite IC-Index scores. T-tests were used to compare means between groups. **(D-E)** Confusion matrices of somatic and germline IC-Indices (cutoff >=5) vs TIME score (cutoff=median).

**Fig S6.**
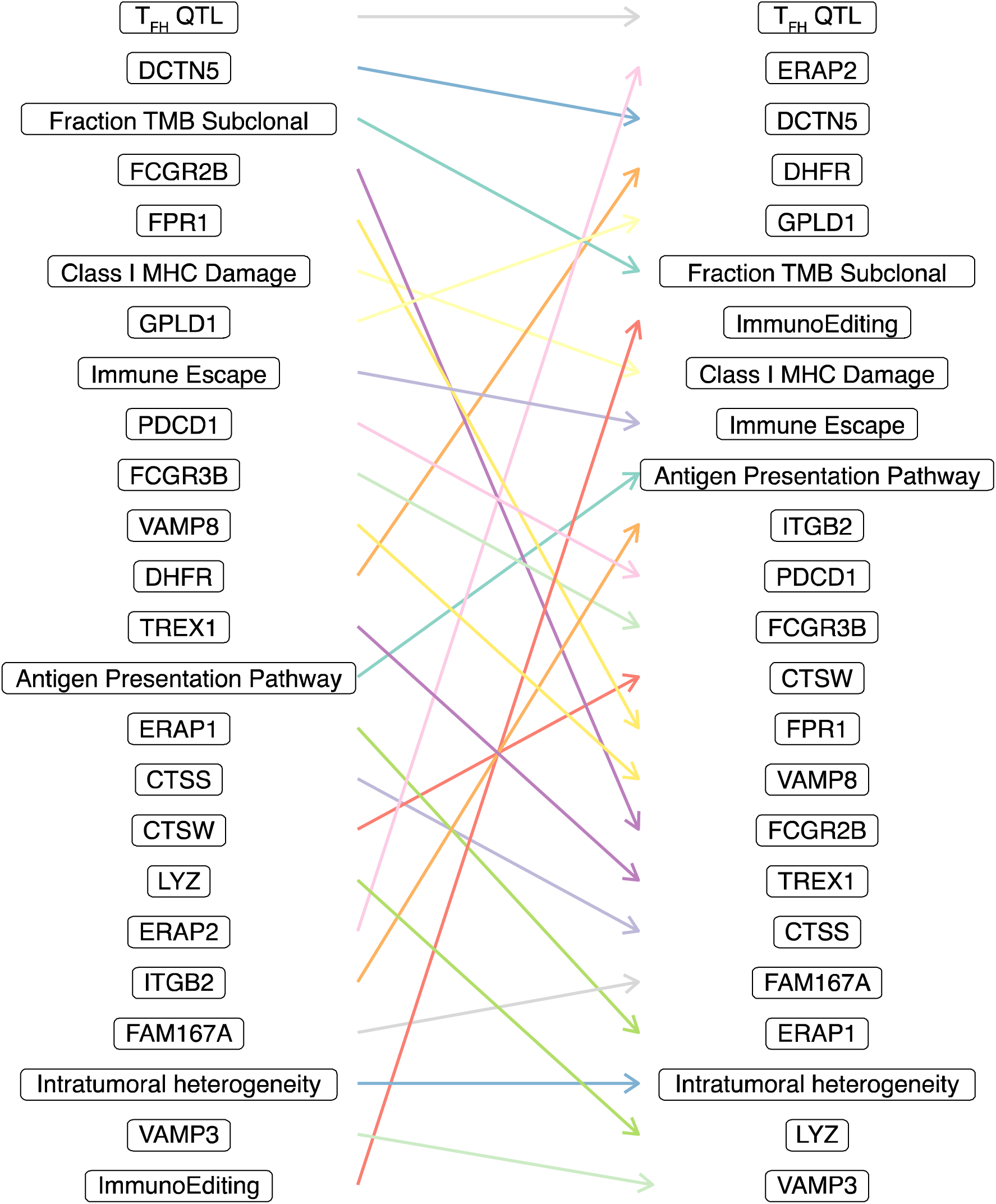
Difference in feature importance rankings of composite model features in linear (left) versus nonlinear (right) models.

**Fig S7.**
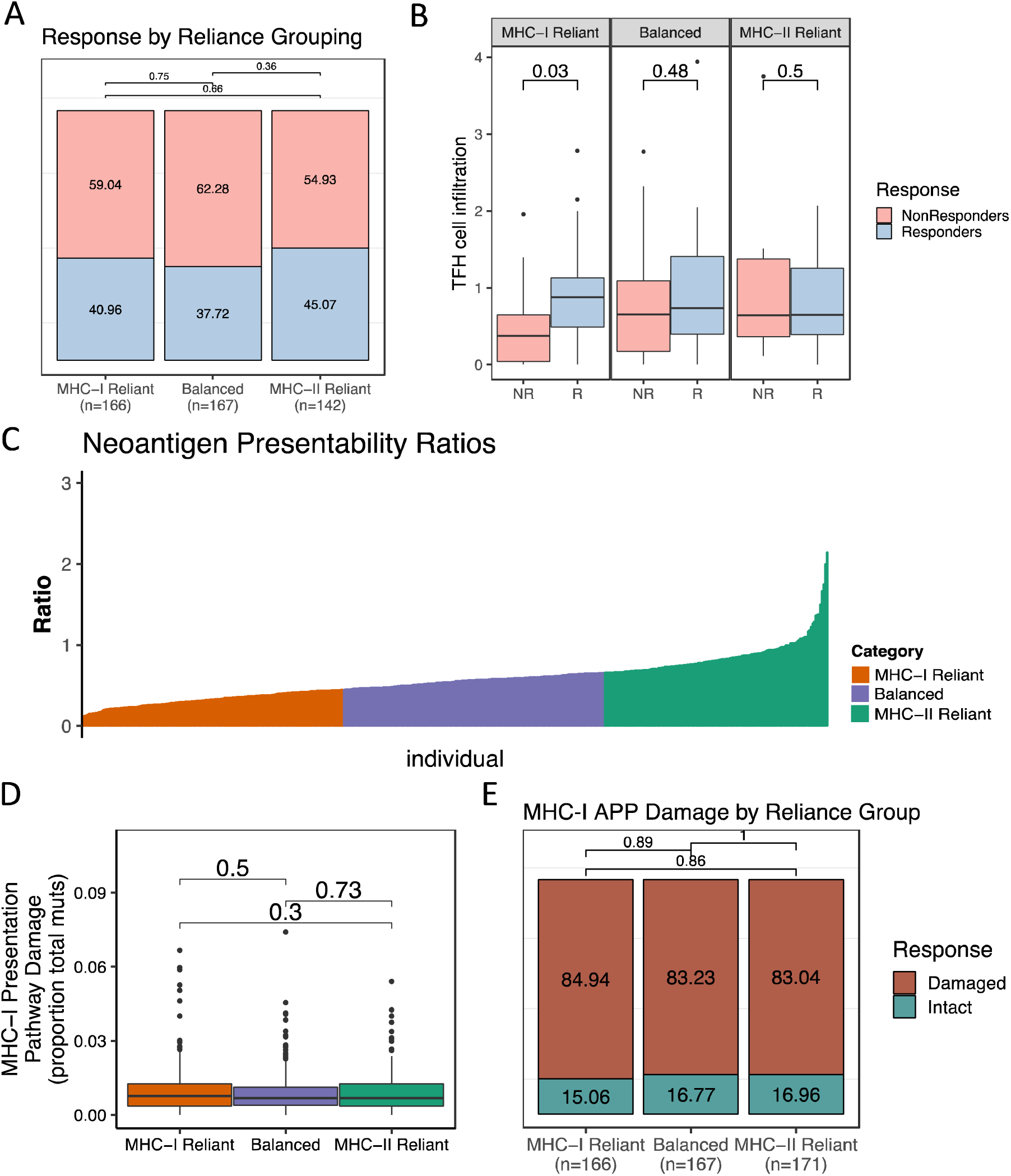
**A.** Response rates by reliance groupings across discovery patients. Chi squared tests were performed between groups. **B.** T_FH_ cell infiltration estimates stratified by MHC-reliance grouping and ICB response. Mann-Whitney U tests were used to compare means between groups. **C.** Waterfall plot of ratio of well-presented MHC-II to MHC-I neoantigens across discovery patients. **D.** MHC-I presentation pathway damage as a proportion of total number of mutations split by MHC Reliance grouping. T-tests were used to compare groups. **E.** Proportion of patients with any damage to MHC-I antigen presentation pathway split by MHC Reliance grouping. Chi squared tests were performed between groups.

**Fig S8.**
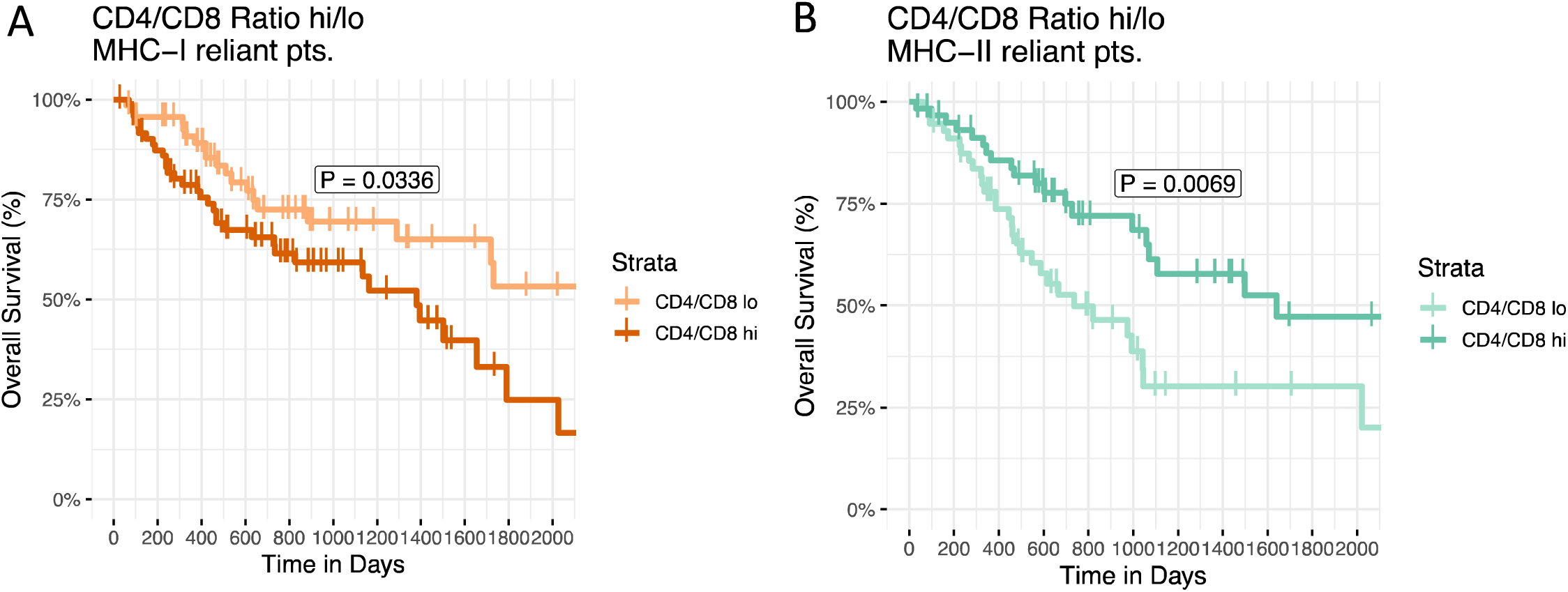
**A.** MHC-I reliant patients in TCGA with melanoma, non-small cell lung cancer or renal cell carcinoma stratified by CD4/CD8 T Cell infiltration ratio. **B.** MHC-II reliant patients in TCGA with melanoma, non-small cell lung cancer or renal cell carcinoma stratified by CD4/CD8 T Cell infiltration ratio. P Values generated via log-rank test

**Fig S9.**
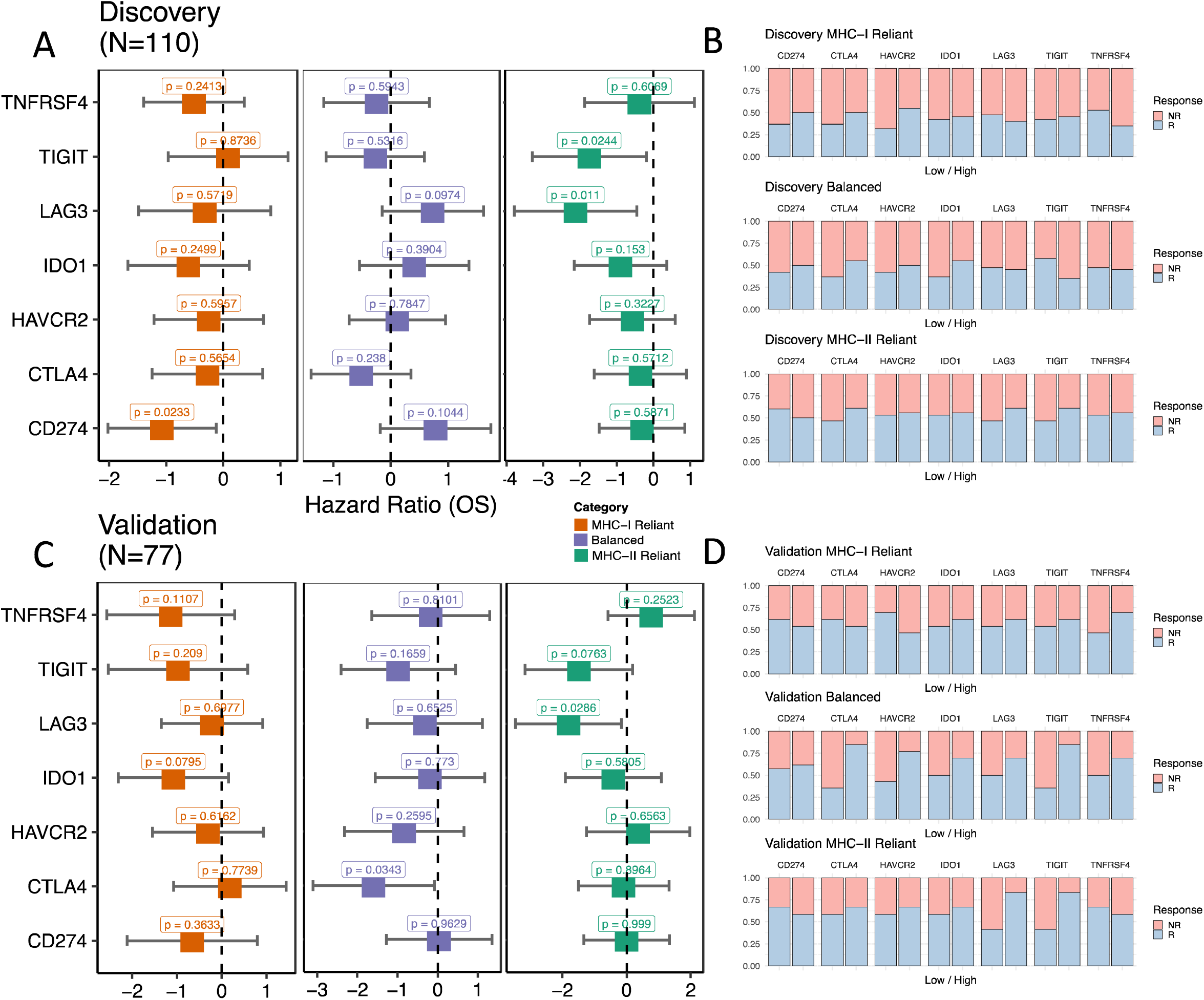
**A.** Univariate analysis of potential checkpoint inhibitors in the discovery samples. **B.** Proportion of patients responding in each MHC reliance group when discovery samples are partitioned according to low versus high expression of 7 different checkpoint genes. **C.** Univariate analysis of potential checkpoint inhibitors in the validation samples. **D.** Proportion of patients responding in each MHC reliance group when validation samples are partitioned according to low versus high expression of 7 different checkpoint genes

**Fig S10.**
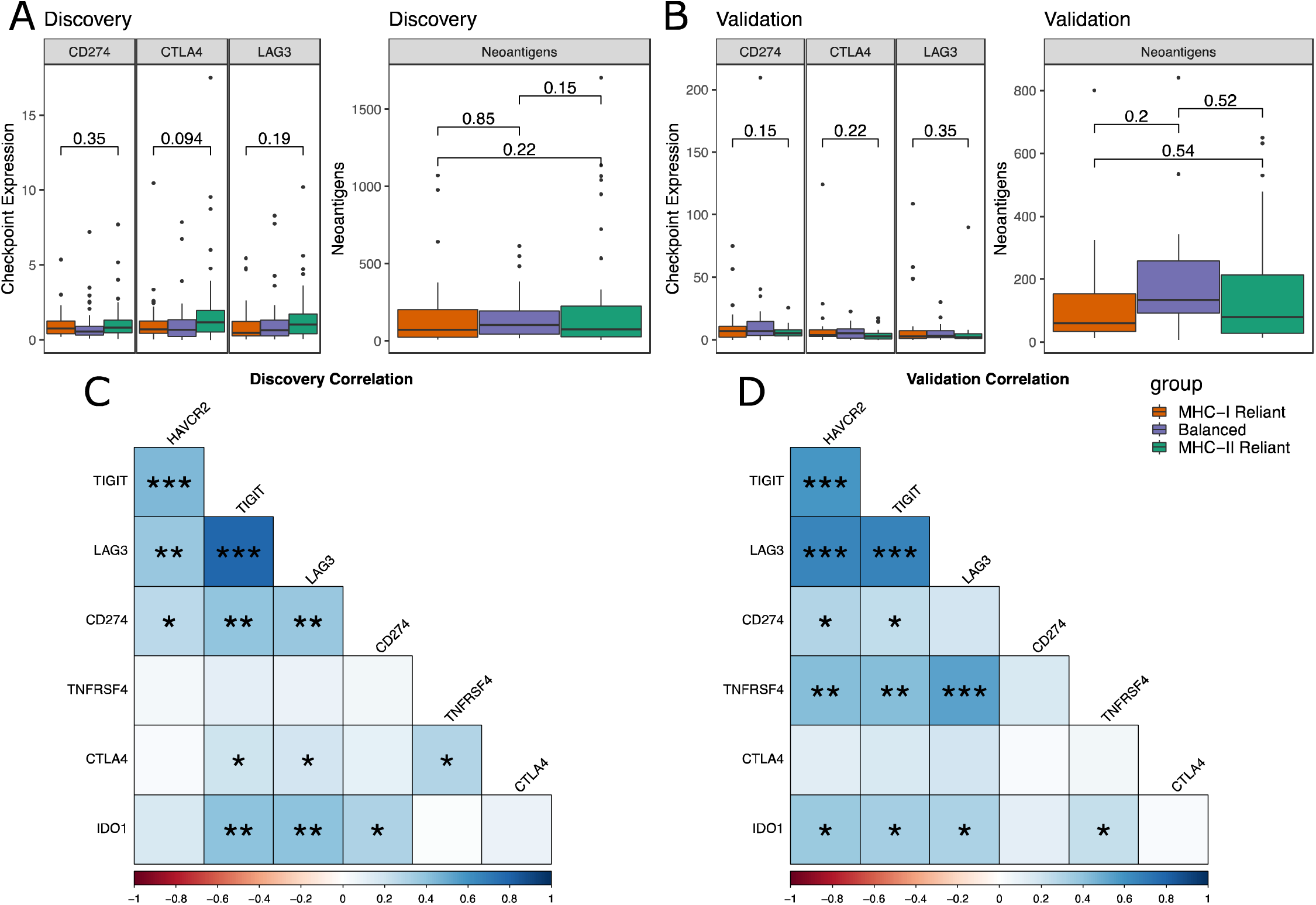
**A.** Checkpoint expression and neoantigen levels by MHC Reliance category in discovery samples. **B.** Checkpoint expression and neoantigen levels by MHC Reliance category in validation samples. **C.** Pearson correlation between various checkpoint genes across discovery samples. **D.** Pearson correlation between various checkpoint genes across validation samples.

**Fig S11.**
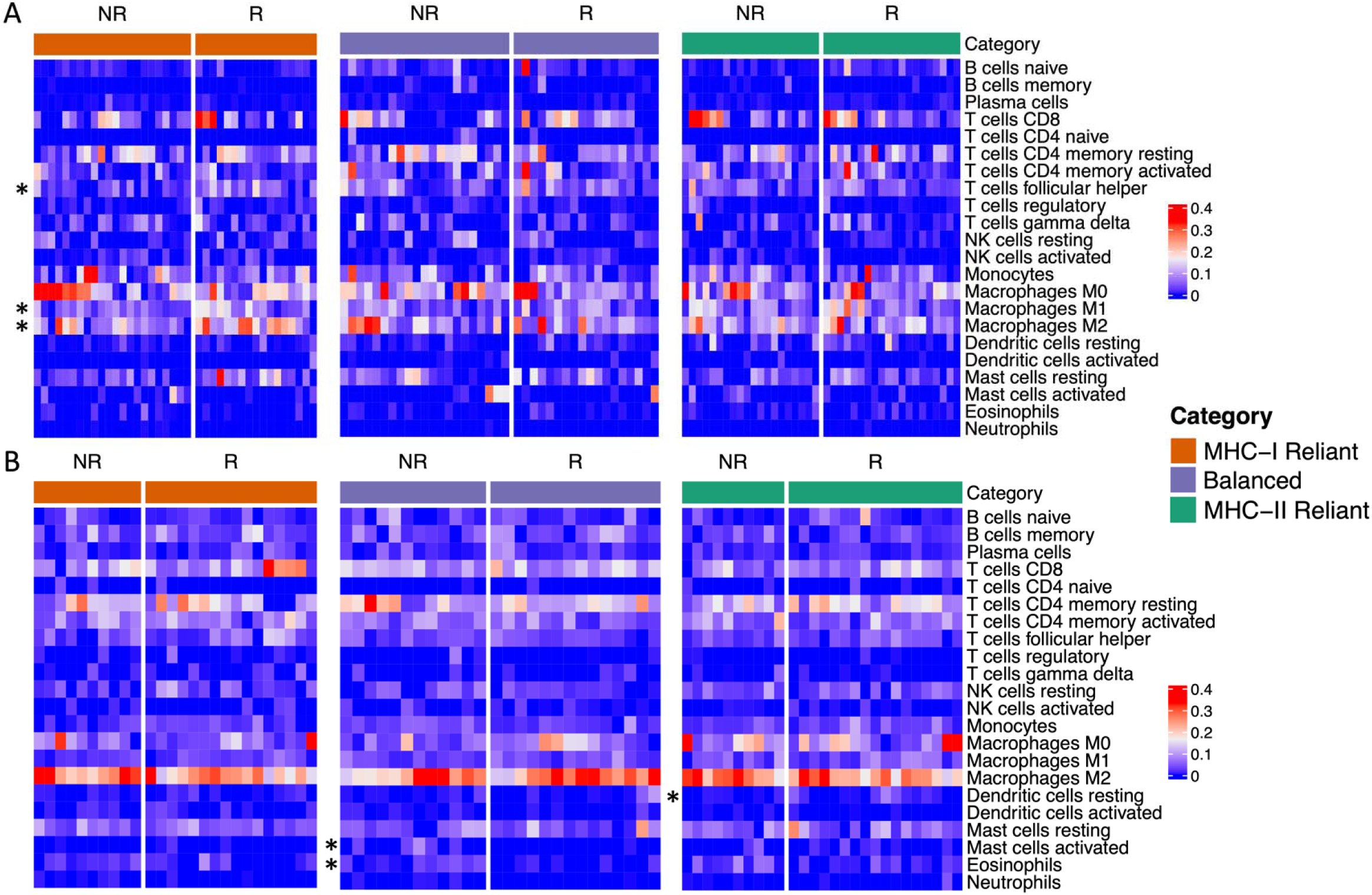
**A-B.** Heatmap of CIBERSORTx derived immune infiltration estimates for discovery **(A)**, and validation **(B)** samples split by response and MHC Reliance category. Mann-Whitney U tests were performed between responders and nonresponders of each MHC Reliance category. P value of <=0.05 are marked with an asterisk.

